# SARS-CoV-2 requires acidic pH to infect cells

**DOI:** 10.1101/2022.06.09.495472

**Authors:** Alex J.B. Kreutzberger, Anwesha Sanyal, Anand Saminathan, Louis-Marie Bloyet, Spencer Stumpf, Zhuoming Liu, Ravi Ojha, Markku T. Patjas, Ahmed Geneid, Gustavo Scanavachi, Catherine A. Doyle, Elliott Somerville, Ricardo Bango Da Cunha Correira, Giuseppe Di Caprio, Sanna Toppila-Salmi, Antti Mäkitie, Volker Kiessling, Olli Vapalahti, Sean P.J. Whelan, Giuseppe Balistreri, Tom Kirchhausen

**Affiliations:** Department of Cell Biology, Harvard Medical School, 200 Longwood Av, Boston, MA 02115, USA; Program in Cellular and Molecular Medicine, Boston Children’s Hospital, 200 Longwood Av, Boston, MA 02115, USA; Department of Molecular Microbiology, Washington University in Saint Louis, 660 South Euclid Avenue, Saint Louis, MI 63110, USA; Department of Virology, Faculty of Medicine, University of Helsinki, Helsinki, Finland; Department of Otorhinolaryngology and Phoniatrics - Head and Neck Surgery, University of Helsinki and Helsinki University Hospital, Helsinki, Finland.; Department of Pharmacology, University of Virginia, Charlottesville, VA 22903; Department of Pediatrics, Harvard Medical School, 200 Longwood Ave, Boston, MA 02115, USA; Department of Allergy, University of Helsinki and Helsinki University Hospital, Helsinki, Finland; Center for Membrane and Cell Physiology and Department of Molecular Physiology and Biological Physics, University of Virginia, Charlottesville, VA 22903; Department of Veterinary Biosciences, University of Helsinki, Helsinki, Finland; Virology and Immunology, Helsinki University Hospital Diagnostic Center (HUSLAB), Helsinki, Finland; The Queensland Brain Institute, University of Queensland, Brisbane, Australia

## Abstract

SARS-CoV-2 cell entry starts with membrane attachment and ends with spike-protein (S) catalyzed membrane fusion depending on two cleavage steps, one usually by furin in producing cells and the second by TMPRSS2 on target cells. Endosomal cathepsins can carry out both. Using real-time 3D single virion tracking, we show fusion and genome penetration requires virion exposure to an acidic milieu of pH 6.2-6.8, even when furin and TMPRSS2 cleavages have occurred. We detect the sequential steps of S1-fragment dissociation, fusion, and content release from the cell surface in TMPRRS2 overexpressing cells only when exposed to acidic pH. We define a key role of an acidic environment for successful infection, found in endosomal compartments and at the surface of TMPRSS2 expressing cells in the acidic milieu of the nasal cavity.

**Significance Statement:** Infection by SARS-CoV-2 depends upon the S large spike protein decorating the virions and is responsible for receptor engagement and subsequent fusion of viral and cellular membranes allowing release of virion contents into the cell. Using new single particle imaging tools, to visualize and track the successive steps from virion attachment to fusion, combined with chemical and genetic perturbations of the cells, we provide the first direct evidence for the cellular uptake routes of productive infection in multiple cell types and their dependence on proteolysis of S by cell surface or endosomal proteases. We show that fusion and content release always require the acidic environment from endosomes, preceded by liberation of the S1 fragment which depends on ACE2 receptor engagement.

**One sentence summary:** Detailed molecular snapshots of the productive infectious entry pathway of SARS-CoV-2 into cells

## Main Text

SARS-CoV-2 cell entry begins with engagement at the cell-surface and ends with deposition of the viral contents into the cytosol by membrane fusion. The first step is binding of the viral spike protein (S) with its cellular receptor, angiotensin converting enzyme (ACE2) (*1–4*). The last step delivers the viral genomic RNA in association with the nucleocapsid protein (N), which is removed for translation of the input genome (*5, 6*). Proteolytic activation of S by additional host-cell factors is necessary for it to function as a fusogen. Cleavage of S by furin in producer cells (*7*) generates the S1 receptor binding subunit non-covalently associated with the S2 fusion subunit. The S protein is cleaved by cell surface or endosomal proteases during virion entry into host cells, that activate the viral fusion machinery (*1, 8–10*). This entry associated proteolysis of S has led to the current model of two routes of infectious cell entry: fusion of viral and cellular membranes at the host-cell surface or fusion following endosomal uptake (*6*).

The cellular proteases that are involved in processing S during entry include the transmembrane serine proteases TMPRSS2 or TMPRSS4 found at the cell surface (*1, 8*), and the endosomal cathepsins that require the acidic milieu of the compartments in which they are enriched (*1, 10*). Processing of S by TMPRSS proteases or by cathepsins, at a site designated S2’, depends on prior cleavage at the furin site in the producer cells (*7, 11, 12*). TMPRSS cleavage has been thought to result in infection from the plasma membrane and cathepsin cleavage, in cells lacking TMPRSS activity, with infection from endosomes (*5, 6*). Chemical inhibitors of TMPRSS or cathepsin proteases in cells in culture indeed show that infection of some cell types is more sensitive to inhibition of endosomal cathepsins whereas others are more sensitive to inhibition of TMPRSS proteases (*1, 9*). TMPRSS inhibitors such as camostat and nafamostat are in clinical development as SARS-CoV-2 therapeutics, further highlighting the need to understand how entry pathways depend upon specific proteases.

To analyze the routes of cellular uptake that lead to successful fusion and release of virion contents into the cytoplasm, we developed a set of new tools that allow direct visual tracking of the uptake of single virions and release of their contents. Using a chimeric vesicular stomatitis virus (VSV) in which SARS-CoV-2 S has replaced the endogenous glycoprotein gene (G), we modified the virus to permit the separate tracking of the S protein and the viral contents. We engineered a structural component of the replicative core of VSV, the phosphoprotein (P), to append to its amino terminus enhanced green fluorescent protein (eGFP), for tracking the virion content. This eGFP-P chimeric virus is structurally fluorescent and depends upon S for entry. We also sparsely labelled the S protein by direct conjugation with a fluorescent dye, allowing us to visualize the steps of entry, from S1 fragment release to membrane fusion, including virion content release into cells during productive infection. We have found that content release requires acidic pH and that it occurs principally from endosomes irrespective of the cell type and irrespective of the dependence of the virus on TMPRSS2 or cathepsin-mediated processing of S. Only mild acidification of the medium allows efficient entry at the plasma membrane. We correlate our findings with studies of infection by several human isolates of SARS-CoV-2.

## RESULTS

### Cell entry of individual virions mediated by SARS-CoV-2 S

We used live-cell fluorescence microscopy to monitor uptake of single VSV-SARS-CoV-2 virions that depend upon SARS-CoV-2 S for infection (Wuhan Hu-1 isolate and Delta and Omicron variants, chimeras generated as described: see Methods and (*13*)) and for the Wuhan variant also containing a fluorescent internal structural protein, P, fused at its amino terminus to eGFP (eGFP-P) (Fig. S1). We counted the number of tagged particles internalized 1 hr post inoculation at MOI 0.5 and found 50-70 particles in Caco-2, Calu-3, and Vero cells, but about three times that number in Vero-TMPRSS2 cells, which overexpress TMPRSS2 (Fig. S1D, E), presumably because of more efficient TMPRSS2 cleavage in the high expression cell line. Examination of the labeled virions by spinning-disc confocal microscopy showed distinct, diffraction-limited puncta with a single, Gaussian distribution of fluorescent intensities (Fig. S2A), consistent with the presence of single virus particles and absence of virion aggregates. We also labeled the S protein sparsely with the fluorescent dye Atto565 NHS ester (25-35 dyes per virion), with minimal effect on particle infectivity (Fig. S2A-D). The double labeling allowed us to track the viral S protein separately from the virion contents.

### SARS-CoV-2 S-mediated entry requires endocytic uptake

We used live-cell volumetric, lattice-light-sheet fluorescence microscopy (LLSM) (*14*) to obtain 3D views of virion cell-entry over time during their uptake into cells (Figs. 1A-E, Fig S3). We incubated cells for 8 min, transferred them to the LLSM, and recorded sequential 3-D stacks from a single cell, acquiring a full stack every 2 sec, for 10 min (300 stacks total). We then moved quickly to an adjacent cell and recorded a similar 10 min series of 300 stacks, after which we moved finally to a third neighboring cell, for 10 min. The sequence of representative 10-plane projections in Fig. 1B shows that particles attached to the cell surface during the initial 8 min incubation, followed by efficient internalization at later time points. The figure shows images from Vero-TMPRSS2 cells; we obtained similar results for Vero, Caco-2 and Calu-3 cells, as shown in Fig S3. The views from 20- and 30-minutes post-inoculation also showed many examples of intracellular spots labeled with eGFP-P only, representing delivery of the VSV ribonucleoprotein core (RNP) into cells visualized as the separation of fluorescent eGFP-P from the membrane-bound, Atto565 labelled S glycoprotein.

**Figure 1.**
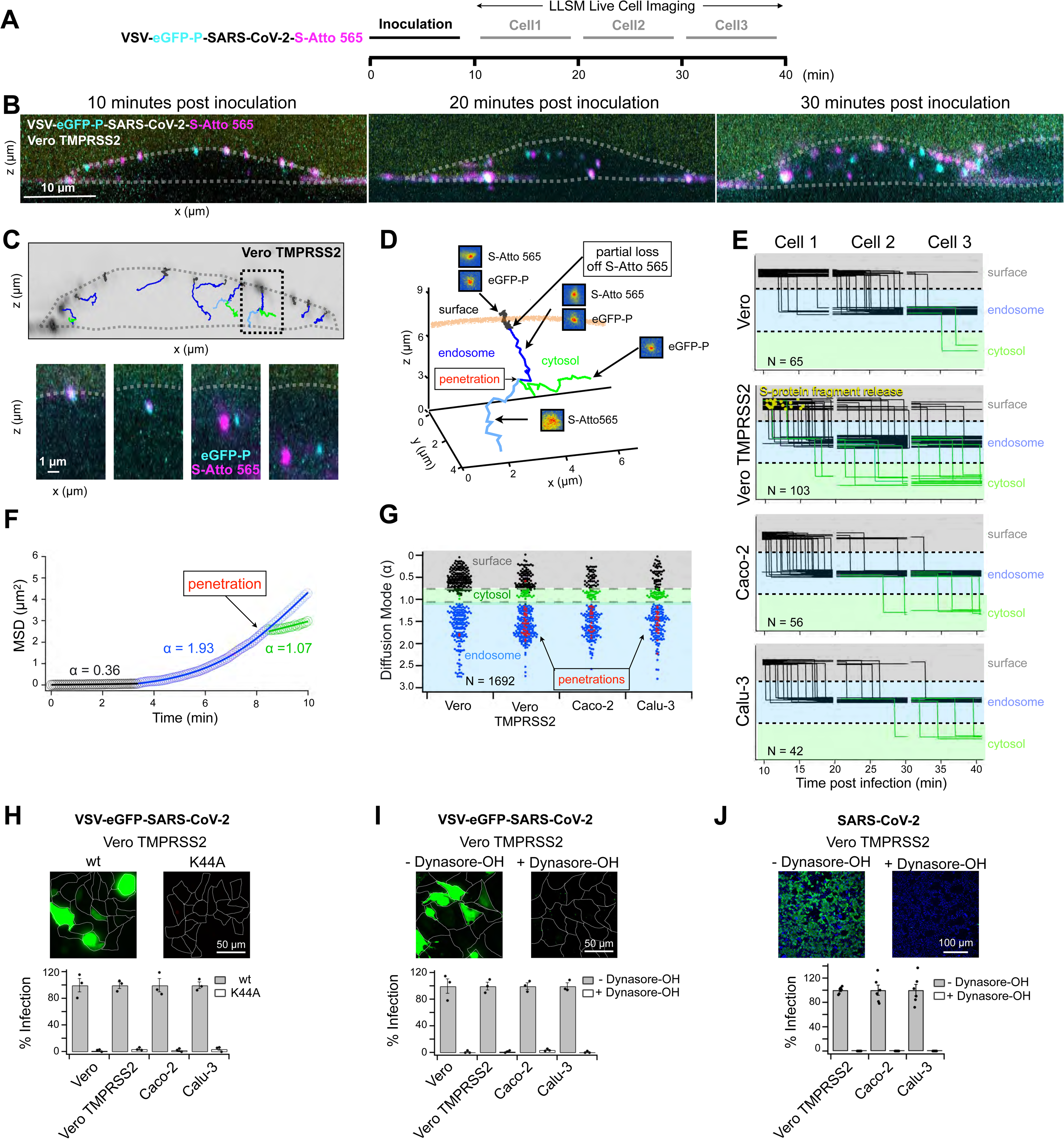
SARS-CoV-2 infection requires endocytosis. **(A)** Schematic of live-cell volumetric lattice-light-sheet fluorescence microscopy (LLSM) imaging experiments **(B-G)** used to obtain 3D time series of VSV-eGFP-P-SARS-CoV-2-S-Atto 565 entry into VERO TMPRSS2 cells during early stages of infection using an MOI of 2. For each experiment, three cells were consecutively imaged volumetrically every 4.7 seconds for 10 min. **(B)** Maximum intensity projections showing fluorescently tagged VSV-SARS-CoV-2 within 1 um in thickness optical sections from the first frame of the time series acquired for representative cells 1, 2 and 3. **(C)** Representative single virion trajectories of VSV-eGFP-P-SARS-CoV-2-S-Atto 565 within a 1 um optical slice of a time series acquired during the first 10-minutes of cell 1. Traces highlight particles at the cell surface (black) and within the cell volume after endocytosis (blue), in both cases containing colocalized eGFP-P and S-Atto 565; it also shows traces in the cytosol (green) containing eGFP-P upon its separation from the Atto565 signal (light blue). Single images highlighting these events are shown in the panels below. **(D)** Orthogonal projection of the traced event highlighted in **(C)**. **(E)** Representative summary of 266 virion traces analyzed during cell entry. Data from single coverslips (out of five) obtained per each cell type are shown. Vertical traces highlight the transfer of virions from the cell surface to the cell interior (assumed to be in endosomes because the colocalization of the eGFP-P with S-Atto 565 signals) or from endosomes to the cytosol (upon loss of localization of the eGFP-P and S-Atto 565 signals). Events corresponding to step wise loss of the S-Atto 565 signal at the cell surface are indicated (yellow). **(F)** Representative plot illustrating the mean squared displacements (MSD) for the trajectory depicted in **(D)** when the particle is at the cell surface (black), in endosomes (blue), or in the cytosol upon separation of eGFP-P (which remains in endosomes, light blue) and S-Atto565 in the cytosol (green). **(G)** Summary dot plot showing the diffusion mode (α) for 1692 virion trajectories and corresponding 139 penetration events; all penetration events occurred from endosomes except for one event at the cell surface in a single Vero TMRSS2 cell. The plot highlights the confined motion (α < 0.80) of virions at the cell surface, trajected motion (α > 1.2) in endosomes, and Brownian motion (0.80< α <1.2) in the cytosol. **(H-J)** Effect of inhibition of endocytosis in the infection by VSV-eGFP-SARS-CoV-2 **(H,I)** or a human isolate of SARS-CoV-2 **(J)**. Top panel shows examples of infection observed in representative fields of Vero TMPRSS2 over expressing or not the dominant negative dynamin K44A mutant or treated or not with 40 µM dynasore-OH. Images in the top panels were obtained using spinning disc confocal microscopy and show maximum intensity projections. Results from similar infections obtained with different cell types are shown in the bottom panel. The difference of results between control conditions and inhibition of endocytosis by K44A dynamin overexpression or incubation with dynasore-OH incubation was statistically significant with p value of <0.001 using an unpaired t-test.

We confirmed that the views in Fig 1B reflected the outcome of sequential internalization and RNP delivery events by tracking individual particles in the volumetric time series acquired using LLSM (Fig. 1C). Single virions attached to cells during the 8 min period following addition of virus, and ∼ 90% (1508/1692) of the attached particles had internalized during the 30 min time course, as recorded in 60 time-lapse 3D videos from 20 cells. The representative 10-plane projection in Fig. 1C obtained during the initial 10-minute interval shows several single-particle examples of virion internalization and three events of RNP core release (for similar results with Vero, Caco-2 and Calu-3 cells, see Fig S3). The highlighted example in Fig. 1C and the complete ortho projection in Fig. 1D show a fluorescent spot corresponding to a virus particle, first captured at the cell surface, then undergoing rapid directed movement towards the cell interior, due to endocytosis and intracellular traffic of virus-containing vesicles. Dissociation of the eGFP-P from the Atto565 signals marks delivery of the genome into the cytosol (see also Figs. S3-S7). In a total of 60 Caco-2, Calu-3, Vero and Vero-TMPRSS2 cells, we detected separation of signals at an intracellular location for 138/1692 trajectories during the 30-min interval starting two minutes after an initial 8-minute inoculation (Fig. 1E), and only one dissociation event at the cell surface (for a Vero-TMPRSS2 cell). The fluorescent signals from eGFP-P released in these times frames remained stable and punctate in the cytosol during the duration of the 10 min acquisition, and any released eGFP-P in the cytosol at the outset of the second or third 10 min time series remained stable for the entire 10 min recording. Moreover, we never observed uncoated (delivered) particles (i.e., released eGFP-P) in the early frames of the time lapse from the first cell. These observations rule out the possibility of rapid, early entry events from the cell surface or from endosomes during the "blind" 8 minutes during inoculation, and we conclude that an endocytic route accounted for all but one of the detected VSV-SARS-CoV-2 fusion events in these experiments.

Analysis of mean square displacement (MSD) curves from three-dimensional, single-particle trajectories allowed us to determine the dynamic regime of the particle at any time point after attachment. The alpha coefficient (α) in the anomalous diffusion equation (<r^2^> = 6Dt^α^) corresponds to confined motion (α < 0.8) for the virus attached to the cell surface, directed motion (α >1.2) for intracellular particles with co-localized Atto565-labelled S and eGFP-P labelled RNP, presumably associated with endosomes, and Brownian motion (0.8< α <1.2) for eGFP-P spots diffusing in the cytosol. The results shown in Fig. 1F (which corresponds to the viral particle traced in Figs. 1 C and D) and the summary for all tracked particles in Fig. 1G (see also Fig. S3-S7) illustrate the moment at which an RNP escapes into the cytosol (α = 1.07), while the tagged S remains associated with an endosome (α = 1.93, see below).

The chimeric VSV particles are roughly 80 by 200 nm (*15, 16*) and appear in the microscope as diffraction-limited spots. Endosomes typically range in diameter from 300-1000 nm (*17*). Thus, a subset of fluorescent endosomes are larger than the diffraction limit, facilitating use of an increased apparent size of an Atto565 fluorescent spot as a proxy for viral membrane fusion with the surrounding (larger) endosomal membrane (Fig. S8). Using this approach, we detected fusion in 17 ± 4 % (n = 20 virions), 28 ± 2% (n = 61 virions), 23 ± 2% (n = 27 virions), and 27 ± 6% (n = 30 virions) of the traces in Vero, Vero-TMPRSS2, Caco-2 or Calu-3 cells, respectively. All the events coincided with release into the cytosol of the eGFP-P labeled RNP core of VSV (Figs. 1E, S3, S7). We did not observe release of the virion contents at the plasma membrane (Figs. 1E, S3-S7), indicating that S-mediated infection of cells only occurred through an endocytic pathway.

### SARS-CoV-2 S-mediated infection requires endocytosis

Dynamin, a large GTPase required for cargo uptake in clathrin-mediated or fast endophilin-mediated endocytosis (*18*), is susceptible to interference by a dominant negative mutant, K44A (*19*). To test whether dynamin-dependent endocytosis is necessary for SARS-CoV-2 S mediated infection, we transiently over expressed the dominant-negative mutant in Caco-2, Calu-3, Vero and Vero-TMPRSS2 cells. We monitored the effect on infection of these cells by VSV-SARS-CoV-2 that expresses eGFP as a marker of infection (VSV-eGFP-SARS-CoV-2) according to the scheme summarized in Fig S9A, and we quantified infection by single-cell imaging at 8 hours post inoculation. In control cells, to achieve comparable infectivity we used 10 times more VSV-SARS-CoV-2 for Caco-2 and Calu-3 cells than for Vero or Vero-TMPRSS2 over-expressing TPMRSS2 (Figs. S9B, C). Irrespective of the cell type, infection was inhibited by the dominant negative dynamin mutant (Fig 1H) or by addition of dynasore-OH, a small molecule dynamin inhibitor (*20*) (Fig. 1I). Dynasore-OH addition at 1h post-inoculation had no effect on viral infectivity, consistent with inhibition of an early, entry-related event, but not with later inhibition of viral gene expression (Fig. S9C). We previously reported that S mediated infection of cells by VSV-SARS-CoV-2 correlates well with infection by SARS-CoV-2 (*9, 13, 21, 22*). In accord with that correlation, dynasore-OH also inhibited infection of Calu-3 and Vero-TMPRSS2 cells by a SARS-CoV-2 Wuhan (B.1) clinical isolate (Fig 1J). These results show a critical role for endocytosis in SARS-CoV-2 infection of multiple cell types in culture including cells that express TMPRSS2.

### Viral entry from endosomes

Our data show that internalized virus particles reached internal membrane compartment(s) from which fusion and genome entry into the cytosol occurred. To establish the identity of the(se) endosomal compartments, we used live-cell LLSM to track particles entering SVG-A cells ectopically expressing ACE2, or both ACE2 and TMPRSS2, and gene-edited for fluorescent labeling of specific endosomal compartments. (Fig. 2A, S10). Single particle tracking of 6815 viruses over a 50 min period starting from 30 minutes post inoculation of cells revealed VSV-SARS-CoV-2 particles trafficked from the cell surface to early endosomes marked with early endosomal antigen 1 fused to the fluorescent protein Scarlett (EEA1-mScarlett) and subsequently to late endosomes/lysosomes marked with a Halo-tagged version of the cholesterol transporter Niemann Pick C1 NPC1-Halo-JFX646 (examples for single virions shown in Fig 2 B, C, S11-S12). We only detected particles located in the interior localizing with the endosomal markers in early frames of the time lapse from the first cell taken at the end of the 30 min inoculation period. Like with the other cells, these observations also rule out for SVG-A cells the possibility of entry events from the cell surface or from endosomes during the 30-min inoculation period.

**Figure 2.**
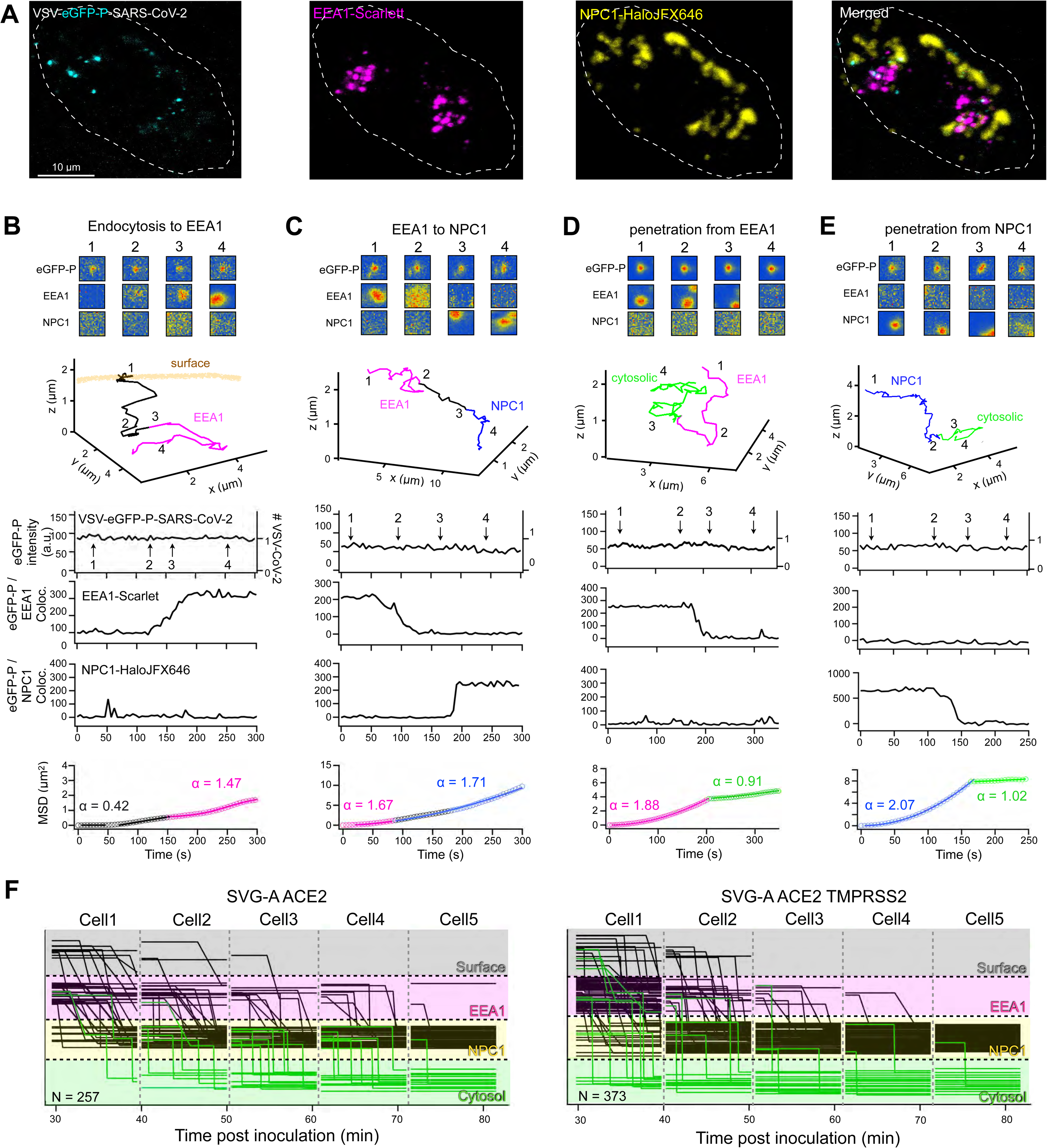
Endocytic entry routes of VSV-SARS-CoV-2. VSV-eGFP-P-SARS-CoV-2 was used to infect SVG-A gene-edited to express early endosomal antigen 1 fused to the fluorescent protein Scarlet (EEA1-Scarlett) as an early endosomal marker and for late endosomal/lysosomal compartments a Halo-tagged version of the cholesterol transporter Niemann Pick C1 (NPC1-Halo) together with ectopic expression of ACE2 and TMPRSS2 and volumetrically imaged using LLSM according to Fig. 1. **(A)** Representative 2 µm projection from the first frame of the time series acquired 8-min after inoculation. **(B-E)** Representative examples of single trajectories of VSV-eGFP-P-SARS-CoV-2 highlighting the extent of colocalization between eGFP-P and EEA1 or between eGFP-P and NPC-1 Halo labeled with JFX646 (top panel), the orthogonal projection of the trajectory (middle panel) and corresponding plots for number of VSV particles on the spot, extent of colocalizations and MSD (bottom panels). Additional examples found in related Fig S11-16. **(I) (F)** Representative summary of results for 257 and 373 virion traces analyzed during cell entry from single coverslips (out of a total of five) plated with SVG-A ACE2 or SVG-A ACE2 TMPRSS2. Vertical and diagonal traces highlight the transfer of virions from the cell surface to its interior and associated with early or late endosomes/lysosomes as defined by colocalization of eGFP-P with EEA1-Scarlett or eGFP-P with NPC1-Halo, respectively.

**Figure 3.**
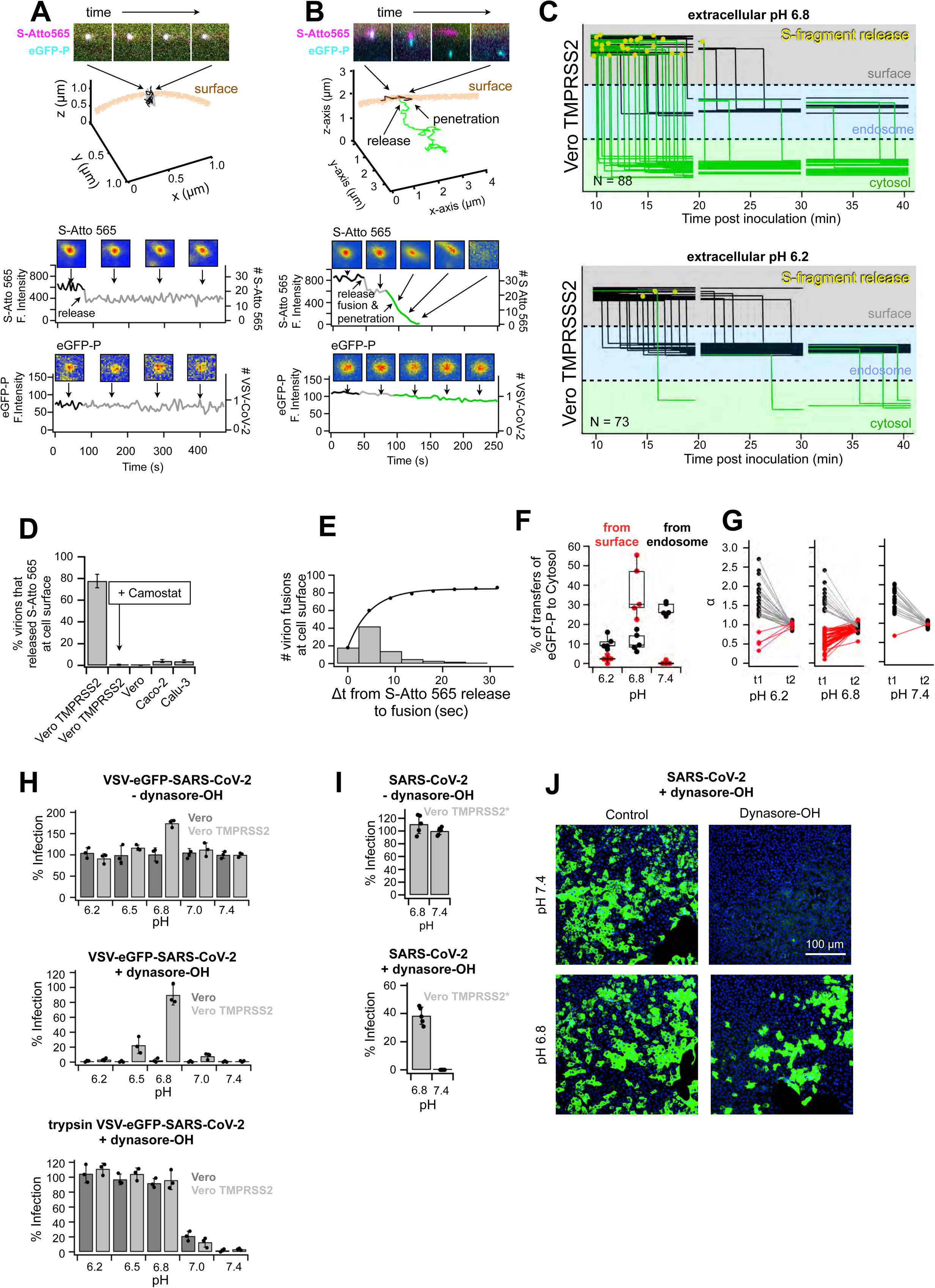
Surface entry route of SARS-CoV-2. **(A-G)** VSV-eGFP-P-SARS-CoV-2-S-Atto 565 was used to infect Vero TMPRSS2 cells and used to study the effect by acidic pH in the medium on the TMPRSS2 mediated surface release of the S-fragment, on cellular location of fusion and genome delivery and on infectivity. (A, B) Single virion trajectories of VSV-eGFP-P-SARS-CoV-2-Atto565 in a Vero TMPRSS2 cell incubated at pH 6.8 showing in (A) an example of S-release at the surface without subsequent fusion and (B) an example of S-release followed by penetration of eGFP-P to the cytosol. Orthogonal views of the tracings and corresponding time-dependent fluorescent intensities for S- Atto 565 and eGFP-P are shown. **(C)** Representative summary from 237 virion traces analyzed during cell entry. Data from single coverslips (out of five) obtained per each pH condition in the medium are shown. Vertical traces of cells incubated at 6.8 highlight the efficient transfer of virions from the cell surface to the cell interior (based on loss of signal colocalization between eGFP-P and S-Atto 565 and corresponding change of diffusion from constrained to directed). Events of stepwise partial loss of S-Atto 565 are indicated (yellow). Similar data with cells incubated at pH 6.2 shows accumulation of virions in endosomes, complete absence of fusion events from the cell surface and limited number of fusion events from endosomes. **(D)** Data showing fraction of virions that released the S-fragment from virions at the cell surface of Vero TMPRSS2 cells incubated at pH 6.8 in the absence or presence of 10 uM Camostat, or of Vero E6, Caco-2 or Calu-3 cells also at pH 6.8 and in the absence of Camostat. The difference of results between control and all other conditions was statistically significant with p value of <0.0001 using an unpaired t-test. **(E)** Cumulative plot corresponding to the dwell time between the stepwise partial drop of the S- Atto 565 signal of a virion at the cell surface and fusion defined by surface spreading of the remaining Atto 565 signal and transfer into the cytosol of eGFP-P. Data from 86 traces and from five experiments. **(F)** Effect of extracellular pH on the transfer of eGFP-P of virions from the cell surface (red) or from endosomes (black) to the cytosol. Each dot represents average +/- std from 5 coverslips with 3 cells imaged per coverslip and at least 600 virus tracked per condition. Line across box represents the median of distribution and the top and bottom represent the quartiles. **(G)** Effect of extracellular pH of the cell medium on the mode of diffusion of the eGFP-P signal associated with a virion before and after delivery from the surface (red) or from endosomes (black) to the cytosol. **(H)** pH bypass infection experiments to test the effect of extracellular acidic pH on the extent of infection of Vero or Vero TMPRSS2 cells by VSV-eGFP-SARS-CoV-2 alone or treated for 30 min with 1 µg/mL trypsin; experiments carried in the absence (top panel) or presence (middle and bottom panels) of 40 uM dynasore-OH. Each data point represents an experiment. In each case, the values determined at pH 6.8 and 7.4 are significantly different with a p value of at least <0.0003 using an unpaired t-test. **(I)** pH bypass infection experiment using authentic SARS-CoV-2 and Vero TMPRSS2* cells in the absence or presence of 40 uM dynasore-OH. Each data point represents an experiment. No statistical difference was observed in the absence of dynasore-OH between pH 6.8 and pH 7.4 (p = 0.13); the difference was statistically significant in the presence of dynasore-OH (p < 0.0001) using an unpaired t-test.

To establish the location of the compartments from which the virion contents were released, we identified the point at which the trajectory of the eGFP-P signal changed from directed (α >1.2) to Brownian (0.8< α <1.2) concomitant with a loss of co-localization with endosomal markers (examples for single virions shown in Fig 2 D, E, S13-S14). All entry events occurred from compartments marked with either EEA1 or NPC1 in cells expressing TMPRSS2, and only from NPC1-compartments in cells lacking TMPRSS2 (Fig. 2 F, S15).

### Release of S1 at the surface of cells expressing TMPRSS2

Membrane fusion mediated by S depends on its proteolytic cleavage and dissociation of the S1 fragment (*6*). Cleavage alone does not release S; the required trigger is ACE2 binding. Addition of soluble ACE2 ectodomain to VSV-SARS-CoV-2 particles activated by trypsin, to bypass TMPRSS2, released about 70% of the Atto 565 label, a reasonable estimate of the fraction of S1 rather than S2. In live-cell imaging experiments with TMPRSS2 expressing cells, we noted a partial loss of Atto 565 intensity from labeled VSV-SARS-CoV-2 particles after they attached to the cell surface (Fig. S19). In these experiments, we used a microfluidics device to introduce virus to cells while in the LLSM, enabling us to detect attachment directly. We interpreted the loss of Atto 565 signal as S1 dissociation, and we took it as a proxy for cleavage by TMPRSS2 at the S2’ site on an ACE2-attached spike protein. Dissociated S1 will remain attached to ACE2 for some time, and in about ∼2 % of the events, we could indeed detect lateral spreading (interpreted as diffusion in the plasma membrane) associated with the abrupt partial reduction of the punctate Atto 565 signal. We found ∼25% loss of the Atto 565 signal per particle in ∼ 78% of particles at the surface of Vero-TMPRSS2 cells expressing TMPRSS2 (Fig. 3 A-D) within the first 10 min of LLSM single-particle tracking (Fig. 1E). In all cases the loss was in a single step with a half time of 2 sec or less (Fig. 3B and Fig. S16). The signal reduction, which preceded endocytosis, was never associated with RNP delivery (e.g., eGFP-P) from the cell surface (Fig. 1E, cell #1), except under the special circumstances of slightly acidic medium during inoculation as described in the following section (Fig. 3 E-J and Fig. 4). The signal loss at the cell surface depended strictly on TMPRSS2 activity, as it was inhibited by 10 µM Camostat and was absent in Vero cells lacking TMPRSS2 (Fig 3D). The decrease in punctate intensity, which we interpret as release of S1, occurred in less than 5% of particles attached to Caco-2 or Calu-3 cells, which naturally express TMPRSS2 (Fig. 3D). This result suggests that in these cells, cleavage at S2’ by TMPRSS2 occurs primarily after uptake into endosomes.

**Figure 4.**
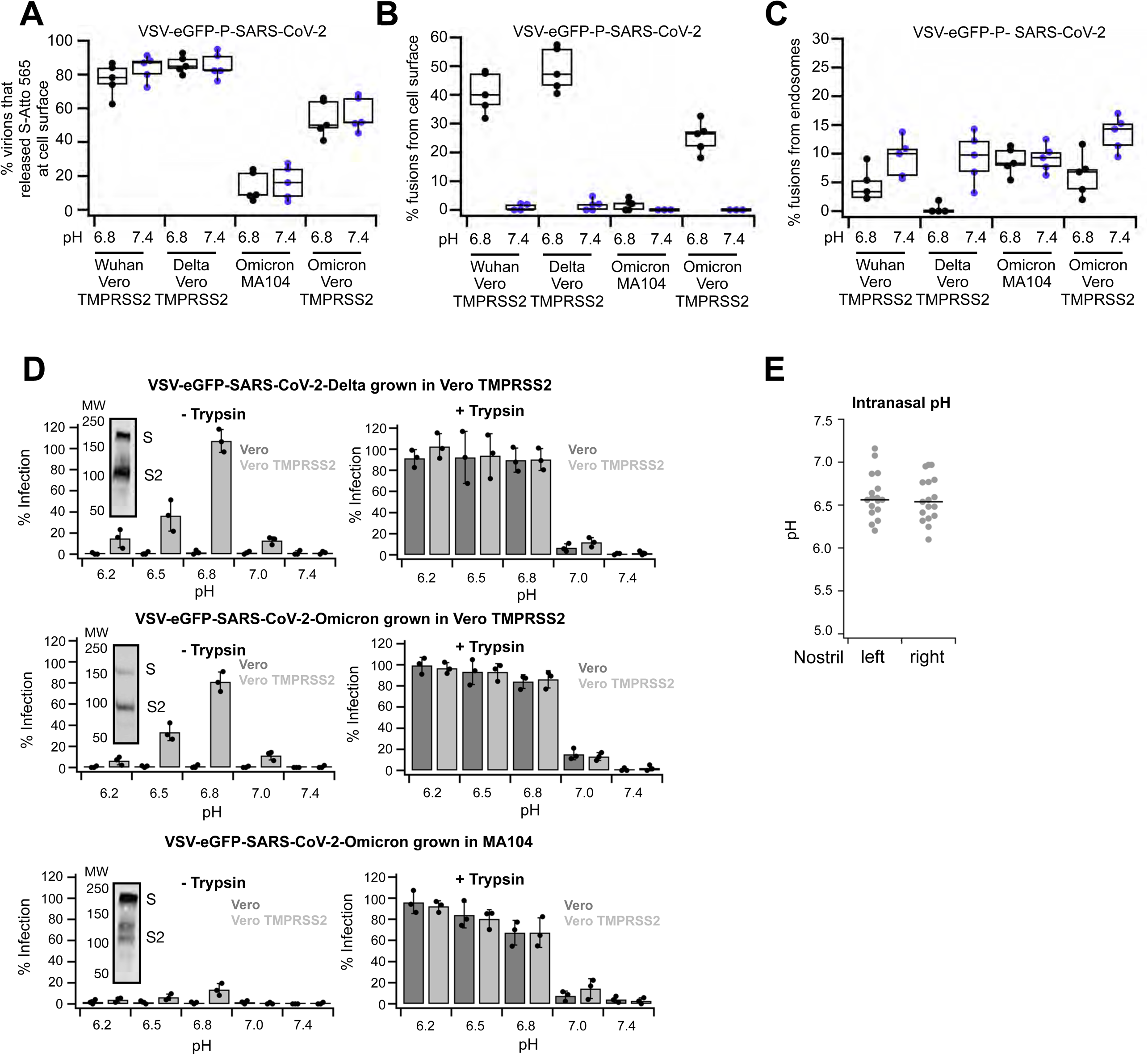
Entry routes of VSV-SARS-CoV-2 variants are conserved. **(A-C)** Effect of extracellular pH and type of cells infected with the indicated variants of VSV-eGFP-P-SARS-CoV-2 on **(A)** the extent of S-fragment release from the cell surface and of **(B)** fusion from the cell surface or **(C)** from endosomes. **(D)** Experiments to determine the effect of extracellular pH on the extent of infection by the Delta and Omicron variants of VSV-SARS-CoV-2 in the presence of 40 uM dynasore-OH. The pH bypass experiments in the right panel were done with trypsin-cleaved virions. The bottom two rows compare results obtained with the Omicron variant grown in MA104 or Vero TMPRSS2 cells. Western Blot showing cleavage states of spike protein of VSV-SARS-CoV-2-Omicron grown in different cell types. The bypass pH experiments in the left panels show statistical differences between pH 6.8 and pH 7.4 (p < 0.0001) for Delta and Omicron grown in Vero TMPRRSS2 and of minimal significance for Omicron grown MA 104 using an unpaired t-test. Similar analysis for the experiments in the right panels show statistical differences between pH 6.8 and pH 7.4 for Delta (p < 0.0001) and Omicron grown in Vero TMPRRSS2 (p < 0.0001) and also for Omicron (P = 0.0015) grown in MA 104. **(E)** Nasal pH values determined from 17 healthy individuals. Each dot represents a single pH determination by the pH catheter at the lower turbinate of the right and left nostrils.

### Viral membrane fusion requires acidic pH

Endosomes undergo rapid acidification, and acidic pH is a trigger that induces fusogenic conformational changes in many viral envelope proteins. We therefore examined the effect of pH on productive viral entry by adjusting the pH of the medium at 10 minutes post inoculation. From 187 traces from VERO-TMPRSS2 cells incubated at pH 6.8, we observed 84 fusion events at the cell surface, as defined by loss of Atto 565 signal and by accompanying eGFP-P delivery into the cytosol (Fig. 3 E-H, S17). The abrupt partial loss of punctate signal, interpreted as S1-fragment release, always preceded fusion (signaled by diffusion in the plasma membrane of the remaining Atto 565 signal with a half-life of about 20 sec) and by eGFP-P delivery into the cytosol, typically within 10 sec (Fig. 3 E, F). We defined eGFP-P delivery into the cytosol by a change in its motion from confined (α <0.8) when on the cell surface to Brownian (0.8< α <1.2) when in the cytosol (Fig. 3G). In the absence of S1-release, we did not detect S1 diffusion or eGFP-P delivery into the cytosol. We conclude that release of S1 and subsequent or concomitant exposure to acidic pH result in release of virion contents into the cell.

Incubation of Vero-TMPRSS2 cells at pH 6.2 rather than 6.8 during inoculation prevented release of the S1-fragment and hence prevented viral fusion at the cell surface (Fig. 3 E, G, and H). This pH dependence is consistent with an expected loss of TMPRSS2 proteolytic activity as the pH falls below 7. When cells were incubated at pH 7.4 throughout infection, we did not observe any particle fusion and content release at the cell surface, even though TMPRSS2 cleavage, as detected by S1-fragment release, still occurred (Fig. 3 E, G and H).

The efficiency of entry by VSV-SARS-CoV-2 tracked by visualizing content release or by extent of infection increased in Vero-TMPRSS2 cells at pH 6.8 (Fig. 3H). To investigate further the requirement of acidic pH for efficient SARS-CoV-2 S-mediated membrane fusion, we carried out bypass experiments to initiate infection directly at the cell surface. We blocked endocytic uptake into cells with dynasore-OH and exposed bound particles to different pH ranging from 6.2-7.4. Infection of Vero, Caco-2 or Calu-3 cells was blocked regardless of the pH, whereas Vero cells overexpressing TMPRSS2 were readily infected by acid triggering of the S protein at the cell surface (Fig. 3J, S18).

These results were consistent with the observed lack of release of the S1-fragment from viruses at the surface of Vero, Caco-2, or Calu-3 cells (Fig. 3D). To eliminate a requirement for TMPRSS2 cleavage of S, we pre-activated particles by incubation with trypsin before inoculating the cells. These pre-activated particles were as infectious in Vero-E6 TM cells treated with dynasore-OH and inoculated at pH < 7 as non-trypsinized particles in the same cells and conditions. Thus, trypsin and TMPRSS2 cleavages were equally effective in activating the particles. With progressive acidification of the medium, infection increased, reaching a maximum at pH 6.6 and remaining constant to pH 6.2 (Fig. 3H). These results show that S-mediated fusion requires pH < 7. We obtained similar results concerning the effect of pH on infectivity of authentic SARS-CoV-2 with cells in which endocytosis was blocked and genome penetration depended solely on cell-surface fusion events (Fig 3I-J).

### Influence of S protein variation on infectious entry pathways

We found that response to infection inhibitors and the requirement for endocytosis and proteolytic cleavage of S for the VSV chimera with the Delta variant S protein were essentially the same for the chimera with the S protein from the original Wuhan-Hu1 isolate (Fig. 4A-D and S20, S21). Conserved events included release of the S1-fragment from receptor-bound virions attached to Vero-TMPRSS2 cells (Fig. 4 B), and S1 release followed by fusion and genome penetration when inoculation was at pH 6.5-6.8 (Fig. 4C). Both Wuhan-Hu1 and Delta also showed enhanced infectivity of Vero-TMPRSS2 cells inoculated at pH 6.8 (Fig. E-G).

Examination of the entry pathway mediated by the S protein of Omicron varied with the cell type used to produce the virus. VSV-Omicron produced in Vero-TMPRSS2 cells depended on TMPRSS2 activation for infection of Vero-TMPRSS2, although somewhat fewer virions released the S1 fragment at the cell surface (Fig 4B-D). VSV-SARS-CoV-2 chimeras with Omicron S produced in MA104 cells could not be activated by TMPRSS2 and failed to infect or fuse from the surface of Vero-TMPRSS2 cells at pH 6.8 (Fig. 4A-D, S20, S21). Infection required activation by cathepsins for fusion and infection from endosomes (Fig 4 A-D, S20, S21). Trypsin treatment of VSV-SARS-CoV-2-Omicron, which cleaves at both the furin and TMPRSS protease sites (*23*), allowed infection to proceed from the cell surface at acidic pH, independent of the producer cell line, even in the presence of an endocytosis inhibitor (Fig.4A-D, S20, S21).

Sequence analysis of the genomic RNA from the VSV-SARS-CoV-2 chimeric viruses confirmed the presence of the TMPRSS2 and furin cleavage sites in Omicron S. As furin cleavage in producing cells is essential for subsequent proteolysis by TMPRSS2 in the infecting cell, we tested whether incomplete furin cleavage explained the differential susceptibility to TMPRSS2 activation. We found much less cleavage of Omicron S for virus produced in MA104 cells (Fig 4D) than for virus produced in Vero-TMPRSS2 cells (Fig. 4D). Particles with furin cleaved S were susceptible to TMPRSS2 activation (Fig 4A-D), released the S1-fragment (Fig 4A), and fused at pH 6.8 (Fig. 4 B,D).

### Intranasal pH

Our results suggest that an acidic environment is required for successful infection, found in endosomal compartments and at the surface of TMPRSS2 expressing cells purposely exposed to mildly acidic extracellular pH conditions. Since expression of TMPRSS2 appears to be highly expressed in a subset of cells located in the nasopharyngeal cavity (*24, 25*), we asked whether this milieu would be acidic. Using a pH catheter placed in the left and right nasal cavities of 17 healthy male and female volunteers, we found a mildly acidic pH of around 6.6 (Fig. 4 E), in agreement with earlier measurements (*26, 27*).

## DISCUSSION

We examined how SARS-CoV-2 Wuhan and its variants Delta and Omicron enter host cells using a combination of high-resolution live cell 3D imaging and quantitative assays for viral infectivity. Real-time tracking of single VSV-SARS-CoV-2 chimeric particles by lattice light sheet microscopy allowed us to visualize directly, with a sensitivity and time resolution substantially greater than any previous work, the key early steps of viral infection: virion binding, release of the S1-fragment upon cleavage by TMPRSS2, viral membrane fusion, and genome penetration into the cytosol. These observations revealed a previously unsuspected requirement to fuse by exposure to pH between 6.5 to 6.8, even after priming by TMPRSS2 and the attendant release of S1 and the S2’ fragment (Figure 5).

**Figure 5.**
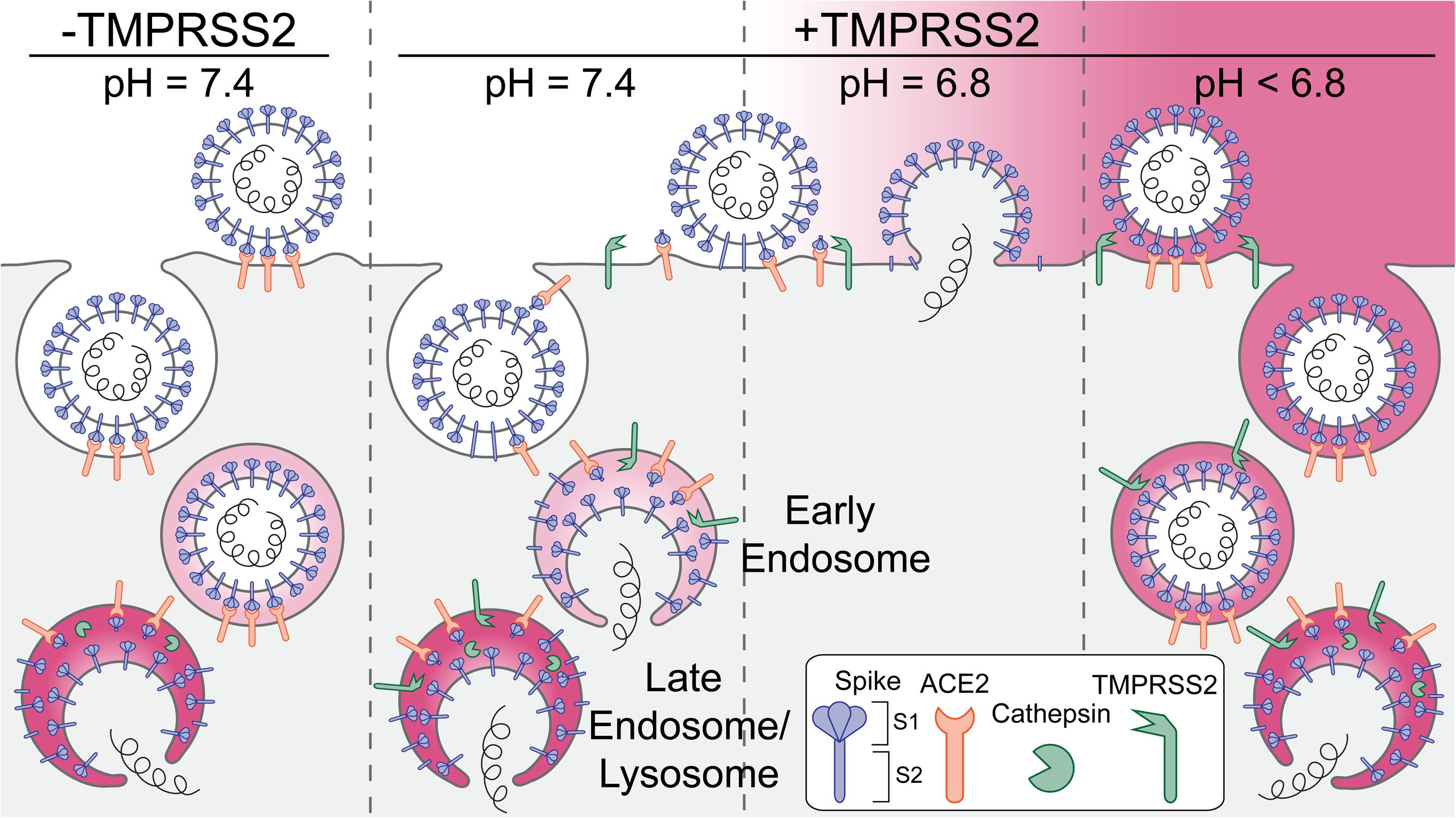
Schematic representation of the principal entry routes SARS-CoV-2 uses for infection. Entry starts with membrane attachment and ends with spike-protein (S) catalyzed membrane fusion releasing the viral contents into the cytosol. Fusion activity depends on two proteolytic cleavage steps, one typically carried out by furin in the producing cell and the second by TMPRSS2 on the cell surface on in endosomes of the target cell. Alternatively, endosomal cathepsins can carry out both cleavages. Exposure of the virus to an acidic milieu is essential for membrane fusion, genome penetration, and productive infection. Fusion and penetration occur only in acidic early and late endosomal/lysosomal compartments but not at the cell surface, even when the furin and TMPRSS2 cleavages have both occurred. Fusion and penetration can occur at the cell surface of cells expressing TMPRSS2 if the extracellular pH is ∼6.8.

Current understanding, derived in part from earlier work on SARS-CoV, has assumed that the only requirement for low pH was for cathepsin L activity when TMPRSS2 was absent or furin cleavage, a prerequisite for TMPRSS2 digestion, had failed during exit from the producing cell (*1, 11, 12, 28–30*). Our results instead distinguish three complementary routes of productive entry, all of which involve exposure of the entering particle to pH < 6.8. The two principal routes are by uptake and traffic to early endosomes, for TMPRSS2-primed virions, or to late endosomal compartments, for cathepsin cleavage in cells regardless of the presence or absence of TMPRSS2. A third, minor route does not require virion uptake, but instead proceeds entirely at the cell surface, but only if cell attachment is at a pH range between 6.5 and 6.8. These observations define the alternative, multi-step pathways of SARS-CoV-2 entry and restrict the conditions for cell-surface penetration.

The released Atto 565-labeled fragment diffused laterally in the membrane, away from the labeled virion, consistent with our identification of the fragment as S1, which we expect to remain attached to ACE2. The release, which always preceded membrane fusion, was independent of exposure to acidic pH. Its dissociation was necessary but not sufficient for S2 to undergo its full, fusion-promoting conformational change. Release required S2’ site cleavage, as evidenced by its absence in the presence of the TMPRSS2 inhibitor, Camostat, and by its absence, even in TMPRSS2 expressing cells, at pH lower than about 6.5, where TMPRSS2 is inactive. Trypsin cleavage in our experiments circumvented TMPRSS2 activity, and we indeed observed fusion at pH as low as 6.2.

The structure of the spike protein and the distribution of lysine residues suggest that most of the Atto 565 label will be on S1 and hence, from the fraction of total label released, that about 25% of the S-protein trimers will have shed S1 upon interaction with membrane bound ACE2, assuming that dissociation of the 3 S1 fragments is cooperative. We estimate from the ratios of stained band intensities on SDS-PAGE that the VSV chimeras have 15-20 S trimers on their surface, and we infer from these numbers that on average, 3-4 spike will have lost S1, liberating their S2 fragments to extend and interact with the host-cell membrane. This estimate is consistent with the likely fraction of the virion surface that makes contact with the cell membrane and with the dependence of S1 shedding from a trimer on its binding to ACE2. It is also consistent with the number of active fusion proteins on other viruses, including VSV itself, required for fusion to proceed (*31–34*).

Syncytium formation between cells expressing SARS-CoV-2 spike and cells expressing ACE2, often used as a spike-mediated fusion assay, does not appear to depend upon acidic pH. But the interface between the two cells will have vastly more spike protein than the interface between a virus and a target cell. Thus, even low probability S1 dissociation events should be sufficient to create a fusion pore, which can widen and spread across the entire cell-cell junction.

Release of S1 from a spike also detaches that spike from ACE2. Because cleavage at the S2’ site was complete in our experiments, any spike bound by ACE2 would probably have released S1. Continued association with the host cell would thus have depended on formation of alternative contacts as S1 dissociates. We propose that formation of an extended intermediate and insertion of the S2 fusion peptides into the host-cell membrane creates the interactions that retain the virion at the cell surface. This proposal further implies that protonation of one or more S2 residues at acidic pH then enables S2 to collapse toward the folded-back, post fusion trimer of hairpins and pull together the viral and host-cell lipid bilayers.

The Omicron variant is more refractory to furin cleavage than previously isolated strains. Its entry pathway will thus depend on the level of furin activity in the producing cells. We indeed found that when grown in Vero-TMPRSS2 cells, Omicron VSV chimeras were susceptible to S2’ cleavage by TMPRSS2 and entered from early endosomes or at the cell surface at mildly acidic pH, but when grown in MA104 cells, they required cathepsin cleavage in late endosomes for entry. With authentic SARS-CoV-2 virus, TMPRSS2 susceptibility similarly depended on the cells in which the virus was propagated. These observations resolve some ambiguities in the literature concerning the role of TMPRSS2 in Omicron infection (*35–37*).

Together with differential protease activities, the pH of respiratory mucosa could also influence viral tropism. The pH of the airway-facing surface of the nasal cavity is between 6.2 and 6.8 (our observations and (*26*). Thus, in principle, rapid entry could occur at the surface of TMPRSS2-expressing cells in the nose. In other parts of the nasopharyngeal cavity and in the lung, the pH is neutral (*38*), and we would not expect virus to fuse in those tissues until its endocytic uptake.

Our suggestion that at neutral pH, a relatively long-lived, extended S2 intermediate may be present at the virus-cell interface bears both on potential therapeutic interventions and on the availability of otherwise occluded epitopes to mucosal antibodies. Persistence of an extended intermediate after gp120 release during HIV entry is thought to account for inhibition of viral infectivity by the peptide fusion inhibitor entfuvirtide and for neutralization by antibodies that recognize epitopes unavailable on a prefusion Env trimer. Our results are consistent with observations that comparable interventions can impede SARS-CoV-2 infection in animal models (*39*).

## MATERIAL AND METHODS

### Materials and Cells

All materials and cells used in this study are described in detail in Supplementary Appendix S1.

### Generation of VSV-SARS-CoV-2 chimeras

The generation of a replication competent recombinant VSV chimera expressing eGFP where the glycoprotein G was replaced with spike (S) protein Wuhan-Hu-1 strain (VSV-eGFP-SARS-CoV-2) has been described (*13*). Additional details for the generation of the VSV recombinants expressing eGFP and SARS-CoV-2 spike variants for Delta (B.1.617.2) and Omicron (B.1.1.529) are described in Supplementary Appendix S1.

### Generation of SVG-A cells expressing ACE2 and TMPRSS2

SVG-A cells ectopically expressing ACE2 and TMPRSS2 were generated by lentivirus transduction. Briefly, lentivirus encoding human ACE2 or TMPRSS2 were generated as follows: HEK293T packaging cells were seeded at 3.8×10^6^ cells in a 10 cm tissue culture plate and grown in complete DMEM supplemented with 10% v/v FBS at 37 °C and 5% CO_2_.

Transfection mixtures containing 90 µL lipofectamine 3000 (Thermo Scientific L3000001), psPAX2 (1.3 pmol; Addgene #12260), pMD2.G (0.72 pmol; Addgene #12259) and TMPRSS2/pLX304 or ACE2/pLJM1 (1.64 pmol; gifts from Sean Whelan) 90 µL lipofectamine 3000 (Thermo Scientific L3000001), 1ul psPAX2 (1.3 pmol; Addgene #12260), 0.6 ul pMD2.G (0.72 pmol; Addgene #12259) and 1.2 ul TMPRSS2/pLX304 or ACE2/pLJM1 (1.64 pmol) in 5 ml OptiMEM medium (Thermo Scientific 31985062) mixed by pipetting and incubated for 20 minutes at room temperature. 0.7 million cells were plated in a 10 cm plate 18 hr prior to transfection; the medium was then replaced with the entire transfection mixture and cells incubated for 6 hours at 37°C, after which the medium was replaced with 15 ml complete DMEM medium. After 12 hours, this medium was replaced with 15 ml of complete DMEM medium, which was then harvested 24 hr later after further growth. After addition of another 15 ml of medium, cells and virus were allowed to growth, ending with a second harvest of medium. The medium was cleared of debris by centrifugation at 5000 x g for 5 mins at room temperature and supernatants containing lentivirus stored at -80°C.

Ectopic stably expression of ACE2 and TMPRSS2 was achieved by transduction of SVG-A cells (gift from Walter J. Atwood, Brown University) gene-edited to simultaneously express fluorescently tagged early and late endosomal markers EEA1 and NPC1 fused to mScarlett or Halo, respectively (*21*). Briefly, SVG-A 1x10^6^ cells were seeded in a well from a 6-well plate and grown overnight in MEM media with 10% FBS. Cleared medium containing ACE2 or TMPRSS2 lentivirus (1 mL) was added to the cells and incubated for 16 h, following replacement with fresh medium cells were incubated for additional 24 hrs. Cells were allowed to grow for another 4 days in the presence of 7 µg/mL puromycin to select for ACE2 expressing cells or 5 µg/mL blasticidin for TMPRSS2 expressing cells. Surviving cells were grown in the absence of puromycin of blasticidin for 4 days and cell stocks frozen and kept in liquid nitrogen. SVG-A cells simultaneously stably expressing ACE2 and TMPRSS2 were obtained by transduction with lentivirus encoding TMPRSS2 and selection with Blasticidin of cells stably expressing ACE2.

### Preparation of VSV chimeras for imaging and infection experiments

All VSV-SARS-CoV-2 variants were grown in MA104 cells in 15 to 20 150-mm dishes and infected at a multiplicity of infection (MOI) of 0.01 as previously described (*9, 13*) in addition to the Omicron variant also grown in Vero TMPRSS2 cells. Briefly, media containing the viruses were collected 72 hours post infection and clarified by centrifugation at 1,000 x g for 10 min at 4°C. A pellet with virus and extracellular particles was obtained by centrifugation in a Ti45 fixed-angle rotor at 72,000 x g (25,000 rpm) for 2 hours at 4°C, then resuspended overnight in 0.5 mL PBS at 4°C. This solution was layered on top of a 15% sucrose-PBS solution and a pellet with virions obtained by centrifugation in a SW55 swinging-bucket rotor at 148,000 x g (35,000 rpm) for 2 hours at 4°C. The resulting pellet was resuspended overnight in 0.4 mL PBS at 4°C, layered on top of a 15 to 45% sucrose-PBS linear gradient and subjected to centrifugation in a SW55 swinging-bucket rotor at 194,000 x g (40,000 rpm) for 1.5 hours at 4°C. The predominant light scattering band located in the lower one-third of the gradient and containing the virions was removed by side puncture of the gradient tube. Approximately 0.3 ml of this solution was mixed with 25 ml of PBS and subjected to centrifugation in a Ti60 fixed-angle rotor at 161,000 x g (40,000 rpm) for 2 hours at 4°C. The final pellet was resuspended overnight in 0.2 - 0.5 mL PBS aliquoted and stored frozen at -80°C for use in subsequent imaging and infection experiments, without detectable change in infectivity.

### Isolation and propagation of SARS-CoV-2

A human isolate of SARS-CoV-2, Wuhan (B.1) was obtained in accordance with the protocol approved by the Helsinki University Hospital laboratory research permit 30 HUS/32/2018§16. Briefly, a nasopharyngeal swabs from a patient infected with COVID19 was suspended in 0.5 ml of universal transport medium (UTM^®^, Copan Diagnostics) and used to inoculate Vero TMPRSS2* cells for 1 h at 37°C, after which the inoculum was replaced with minimum essential medium (MEM) supplemented with 2% heat inactivated FBS, L-glutamine, penicillin, and streptomycin and virus allowed to grow for 48 hr. Virions in the supernatant (P0) were subjected to a similar second round of propagation (P1), analyzed by DNA sequencing, and aliquots stored at 80°C in a solution containing MEM, 2% heat inactivated FCS, 2 mM L-glutamine, and 1% penicillin-streptomycin. Extent of virus replication was determined by real-time PCR (RT-PCR) using primers for SARS-CoV-2 RNA-dependent RNA polymerase (RdRP) (*40*).

### VSV-eGFP-SARS-CoV-2 infection assays

Infection assays for the Wuhan, Delta or Omicron VSV-eGFP-SARS-CoV-2 chimeras were done at a final ∼ 80% confluency of cells plated one day before the infection assay as previously described (*9*) and further explained in Supplementary Appendix S1.

### SARS-CoV-2 infection assays

All experiments with SARS-CoV-2 were performed in biosafety level 3 (BSL3) facilities at the University of Helsinki with appropriate institutional permits. Virus samples were obtained under Helsinki University Hospital laboratory research permit 30 HUS/32/2018§16. Infections were carried for 16 hours at 37°C with 5% CO_2_. Cells were then fixed with 4% paraformaldehyde in PBS for 30 min at room temperature before being processed for immunodetection of viral N protein, automated fluorescence imaging, and image analysis. The detailed protocol is outlined in the Supplementary Appendix S1.

### VSV-SARS-CoV-2 Atto 565 labeling and single molecule Atto 565 dye calibration

Stock solutions of VSV-SARS-CoV-2 and its variants at a concentration of ∼150 µg/ml viral RNA were conjugated with Atto 565-NHS ester (Sigma-Aldrich, cat. 72464) as previously described (*41*). The number of Atto 565 molecules attached to a single virion was determined by comparing the total fluorescence intensities associated with a given virion and the fluorescence intensity associated with the last bleaching step of the same virion, as previously described (*42, 43*). A brief description of these steps are outlined in the Supplementary Appendix S1.

### Trypsin cleavage of VSV-eGFP-SARS-CoV-2

VSV-eGFP-SARS-CoV-2 and variants (as indicated in text) at a concentration 30 µg/mL virus RNA in a total volume of 100 µL in DMEM with 25 mM HEPES, pH 7.4 were incubated for 30 min at 37°C with 1 µg/mL trypsin (Pierce trypsin, TPCK treated from Thermo scientific cat. PI20233). Trypsin activity was terminated with 10 µM Aprotinin (bovine lung, Sigma-Aldrich cat. A1153). The required concentration of trypsin was determined by trypsin serial dilution, aiming for the largest infectivity Vero cells whose endogenous cathepsin proteases activity had been inhibited with 20 µM E-64 (Figure S19). Infections were done with VSV or its variants at a concentration of 0.5 µg/mL (for Vero and Vero TMPRSS2) or 5 µg/mL virus RNA (for Caco-2 or Calu-3). When required, the trypsin cleaved VSV’s were used together with infection inhibitors or for pH bypass experiments as described above.

### *In vitro* release of S1 from trypsin activated VSV-SARS-CoV-2

VSV-eGFP-SARS-CoV-2-Atto 565 was incubated with trypsin at various concentrations for 30 min and the reaction stopped by addition of aprotinin to a final concentration of 10 µM. Virus was then used to infect Vero cells with endogenous cathepsin proteases inhibited with 20 µM E-64 to determine maximum concentration of trypsin required to proteolytically activate the virus. Virus either without or with trypsin cleavage, or with ACE2 bound before and after trypsin cleavage were plated onto a poly-D-lysine coated glass to determine the number of Atto 565 fluorescence dyes associated with each one of the single particles using spinning disc confocal microscopy. At least 8,000 particles were imaged per experimental condition to guarantee the ability to distinguish intensity losses of at least 20%. Every experimental condition was repeated in triplicate and the intensity of the mean intensities of the peak Gaussian fit was used to determine the after and deviation of Atto565 dyes per condition, Figure S19.

A computation simulation was performed to validate the ability to detect with statistical significance a 20% fluorescence intensity loss approximate equivalent to the 20-30% loss observed *in vivo* with virions when attached to the cell surface of Vero TMPRSS2. Experimental data corresponding to 8,845 undigested control virions were used to generate a probability density function from which 9 fit parameters (3 mean intensities with corresponding sigmas and the relative contribution to the distribution) were obtained by fitting the sum of 3 Gaussian distributions. These parameters were then used to generate 8845 random numbers of the same distribution and compared to the experimental data set to illustrate the accuracy of the simulation. A second set of 8845 random numbers were generated with mean intensities reduced by 20%, to generate a probability density distribution representing the fluorescence intensities after ACE2 mediated release of S cleavage by TMPRSS2.

### Nasal pH and temperature measurements

The pH and temperature of the nasal cavity, close to the under lower turbinate, from each nostril from 17 healthy volunteers, age 26-55, males and females, were determined using a Digitrapper pH 400 recorder (Medtronic) connected to a single-use Versaflex disposal dual sensor pH catheter (Medtronic) and a Beurer FT 15/1digital thermometer (Beurer), respectively. The temperature dependent pH response of the Digtrapper was corrected to consider the temperature in the nostrils determined for each volunteer.

These measurements were obtained under the ethical permit n. HUS/2502/2020 granted by the ethical committee of Helsinki and Uusimaa hospital district to Ahmed Geneid and Markku Patjas at the Helsinki University Hospital.

### Preparation of glass coverslips

Infection and uptake assays of VSV-eGFP-SARS-CoV-2 and VSV-P-eGFP-SARS-CoV-2 done by spinning disc confocal microscopy visualization, were performed using 25 mm #1.5 coverslips bound with polydimethylsiloxane (PDMS) of about 1 mm in thickness and of 3 mm (infection) or 5 mm (uptake) in diameter wells punched as previously described (*9*) (see also in Supplementary Appendix S1).

### Live cell spinning disc-confocal microscopy

Visualization experiments were done with an inverted spinning disc confocal microscope (*42*) following the details described in Supplementary Appendix S1.

### Live cell lattice light sheet microscopy

Cells were plated in a 35 mm culture dish containing 5 mm in diameter glass coverslips to achieve 60% final confluency the day of each experiment. Immediately before lattice light sheet visualization, the cover slip was placed on top of parafilm placed in a 10 cm in diameter petri dish including wet chem wipes to maintain humidity. Approximately 10 µL of a solution containing VSV-SARS-CoV-2 in DMEM with 25 mM HEPES at pH 7.4 for the times indicated in the text, after which the coverslips were transferred to the imaging stage of the LLSM. Visualization was done using phenol red free media, (FluoroBrite™) supplemented with 5% FBS and 25 mM HEPES at the indicated pH. Imaging was done at 37°C in the presence of 5% CO_2_ and 100 nM fluorescent Alexa 647 or Alexa 549 dyes added to the medium to determine the cell boundary. The LLSM was operated in sample scan mode with 0.5 µm spacing between each plane along the z-imaging axis and samples imaged as a time series of stacks acquired 1 to 5 sec using dithered multi-Bessel lattice light sheet illumination (*40, 43*). When using gene edited SVG-A cells expressing EEA1-Scarlett and NPC1-Halo, Halo was first labeled by incubation of the cells with 200 nM JFX646 for 30 minutes at 37°C and 5% CO_2_ followed by three 2-minute washes with DMEM containing 10% FBS, 25 mM HEPES, pH 7.4 within 2 hours prior to virus addition to the cover slip. Time series containing 120-300 z-stacks, sequentially obtained every 1-5 sec were acquired with ∼ 10 ms exposures per channel.

The following protocol was used to determine the dwell time between binding of trypsin activated VSV-SARS-CoV-2-Atto 565 Wuhan and release of the S-fragment. Briefly, trypsin activated virions at 5 µg/mL viral RNA were flowed on top of Vero cells plated in a homemade microfluidic flow cell one day prior to the experiment to achieve a 90% confluency. Soluble 100 nM Alexa 549 added to the medium was used to determine the cell outline. Samples were imaged with an AO-LLSM microscope configured for sample scan imaging to acquire every 4 sec a stack with planes separated by 0.6 um (0.3 um along the z-optical axis) using an exposure of 3 ms/plane.

### Single virus tracking and image analysis

The 3D stacks obtained using LLSM were deskewed and the diffraction limited spots were detected and tracked in three dimensions using the automated detection algorithms that uses least-squares minimization numerical fitting with a model of the microscope PSF approximated by a 3D Gaussian function and implemented using the MATLAB developed previously (*40*) available for download (https://github.com/VolkerKirchheim/TrackBrowser_Matlab.git). Estimated fluorescent intensities associated with each spot were calculated from the corresponding amplitudes of the fitted 3D Gaussian and compared to those from single virions bound to poly-D-lysine coated glass imaged under the same acquisition conditions and whose dye content was determined by single bleaching steps (*43*).

Tracks with intensities corresponding to single virus were exported into a custom-made program written in LabView for visualizing trajectories available for download (https://github.com/VolkerKirchheim/TrackBrowser_LabView.git). Each virus trajectory was visually examined for co-localization within a specific compartment (cell surface, EEA1 early endosomes or NPC-1 late endosomes/lysosomes, cytosol) and the mean squared displacements (MSDs) were calculated from all 3D coordinates for all possible time frames within that compartment. A non-linear relationship between MSD and time for anomalous diffusion was fitted to the MSD data according to the power law in equation:

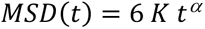

where *K* is the generalized diffusion coefficient and *α* is the anomalous exponent.

### Statistical analysis

An unpaired t-test was used to determine the statistical significance in the difference between control and experimental values.

## ACKNOWLEDGMENTS

We thank Stephen C Harrison for comments, suggestions, and extensive editorial assistance and members of our laboratories for help and encouragement; the staff at HUSLAB Virology and Immunology for providing human nasal swabs for virus isolation, to Suvi Kuivanen and Teemu Smura for virus propagation, sequencing and discussions, and Sanna Mäki for excellent technical work; Elliott Somerville (Kirchhausen laboratory) for excellent laboratory management; Tegy John Vadakkan for maintaining the spinning disc confocal microscope; Lena Tveriakhina (Blacklow laboratory, Harvard Medical School) for western blot analysis.

## Funding

NIH Maximizing Investigators’ Research Award (MIRA) GM130386 (TK)

NIH Grant AI163019 (SPJW, TK)

Danish Technical University (TK)

SANA (TK)

Harvard Virology Program, NIH training Grant T32 AI07245 postdoctoral fellowship (AJBK)

Academy of Finland research grant 318434 (GB, RO)

Academy of Finland research grants 335527 and 336490 (OV)

Helsinki University Hospital Funds TYH2018322 (OV)

University of Helsinki Graduate Program in Microbiology and Biotechnology (RO)

## Author contributions

Alex J.B. Kreutzberger carried all the experiments except those with human SARS-CoV-2 isolates. Louis-Marie Bloyet and Spencer Stumpf generated, characterized, and sequenced the recombinant virus VSV-eGFP-P-SARS-CoV-2 Whuhan. Zhuoming Liu generated, characterized, and sequenced the recombinant viruses VSV-eGFP-SARS-CoV-2 Delta and Omicron. Anwesha Sanyal generated SVGA-A ACE2 and SVGA-A ACE2 TMPRSS2, maintained the cells lines and assisted with infectivity assays. Catherine A. Doyle helped analyzed data and carried out some VSV-SARS-CoV-2 infection assays. Elliott Somerville helped with virus sample preparation for DNA sequence. Gustavo Scanavachi and Anand Saminathan helped collect LLSM imaging data. Giuseppe Di Caprio set up the single particle data analysis pipeline. Volker Kiessling developed the LabVIEW-based viewer, simulated data and produced the single-particle MSD fitting algorithm. Sean P. J. Whelan oversaw the work to generate and characterize the VSV-chimeras, participated in early discussions, and helped edit portions of the manuscript. Giuseppe Balistreri guided the work on SARS-CoV-2 and performed infection assays, imaging, and image analysis. Ravi Ojha performed SARS-CoV-2 infection assays, imaging, and image analysis. Olli Vapalahti coordinated the BSL3 work, provided SARS-CoV-2 and performed virus sequencing.

Tom Kirchhausen and Alex J.B. Kreutzberger were responsible for the overall design of the study; Tom Kirchhausen drafted the manuscript with direct input from Alex J.B. Kreutzberger; all authors commented on the manuscript.

## Data and material availability

All materials and data generated in this study are available upon request. Further information and requests for resources and reagents should be directed to and will be fulfilled by the lead contact, Dr. Tom Kirchhausen, <u>mailto:</u>. Requests for VSV-SARS-CoV-2 chimeras and their materials transfer agreements (MTA) should be directed to and will be fulfilled by Dr. Sean Whelan. This study did not generate any unique large-scale datasets. The LabView code for visualizing trajectories is available for download (GITHUB).

---

**Figure S1.**
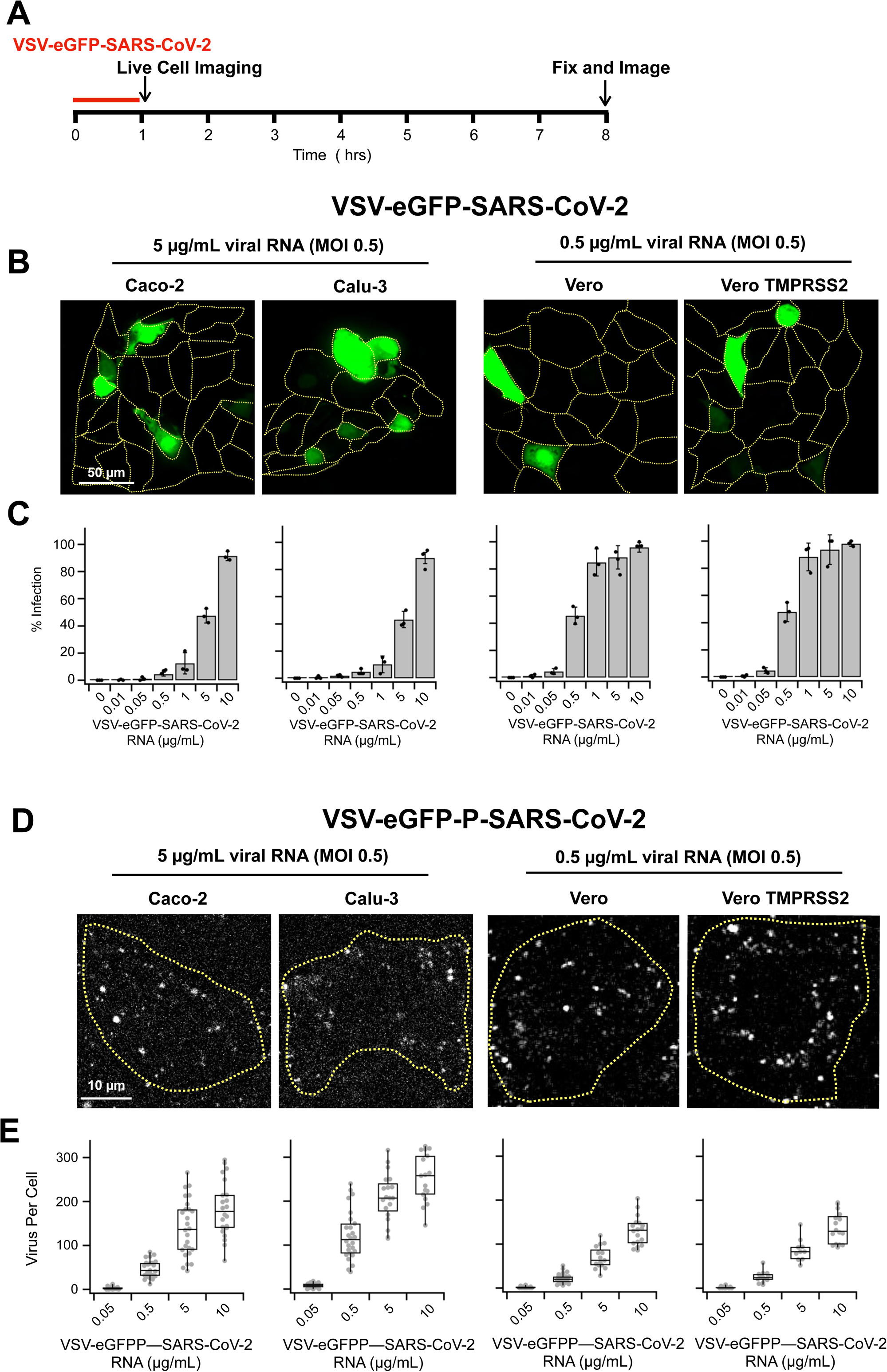
Virus infection and uptake in different cell lines. **(A)** Schematic of infection (fixed cell imaging with spinning disc) or virus uptake (live cell imaging with LLSM) protocols. **(B, C)** Example images **(B)** and quantification **(C)** of VSV-eGFP-SARS-CoV-2 infection in Caco-2 (left), Calu-3 (middle left), Vero (middle right), and Vero TMPRSS2 (right). Each condition contained 3 independent experiments. **(D, E**) Example images and quantification of **(E)** VSV-eGFP-P-SARS-CoV-2 uptake. Each condition contained 3 independent experiments.

**Figure S2.**
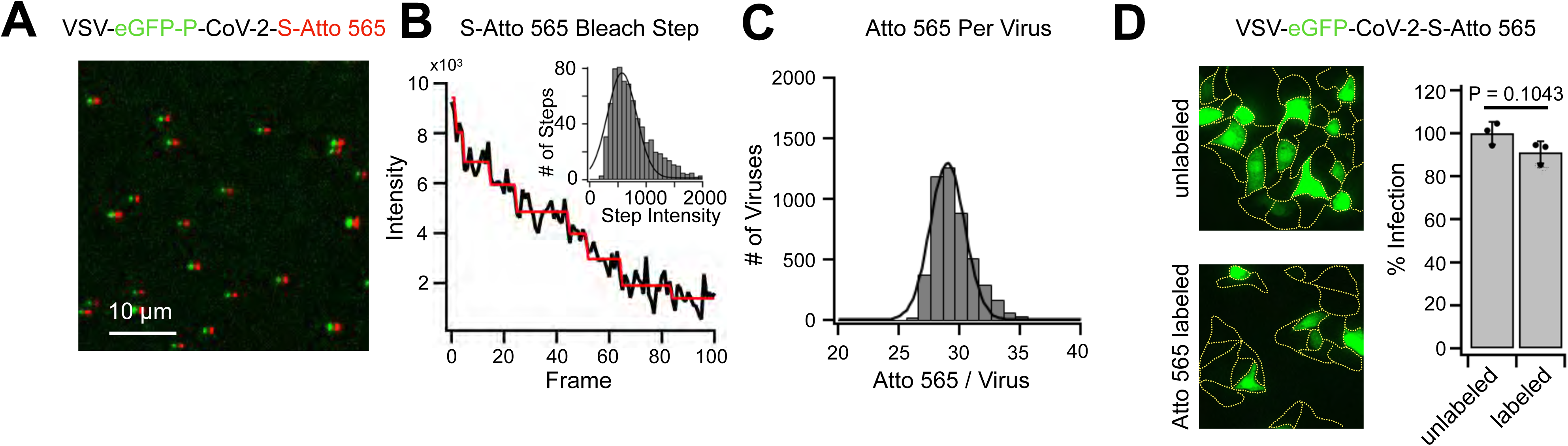
VSV-eGFP-P-SARS-CoV-2-Atto565 labeling. **(A)** Maximum intensity projection image of 5 z-planes spaced 0.27 µm of VSV-eGFP-P-SARS-CoV-2-Atto 565 bound to poly-lysine coated glass collected on a spinning disc-confocal microscope. Shift by 5 pixels of channels highlights colocalization of the fluorescence signals for eGFP-P and S Atto 565 for each virion spot. **(B)** Single molecule photobleaching steps of Atto 565 dye attached to virus taken at 1 second exposure with the step intensity distribution of the last step of 727 VSV-eGFP-P-SARS-CoV-2-Atto565 virus shown in inset. **(C)** The linear relationship between exposure times and fluorescence intensity allows the step size taken at 1 second exposure to be used to calibrate the number of dyes per virus when taken at 20 ms exposures when the dye single on each virus spot is not (*1*). This was used to determine an average distribution of ∼30 dyes per VSV-SARS-CoV-2-Atto 565. **(D)** Labeling VSV-eGFP-SARS-CoV-2 with Atto 565 showed little inhibition of viral infection.

**Figure S3.**
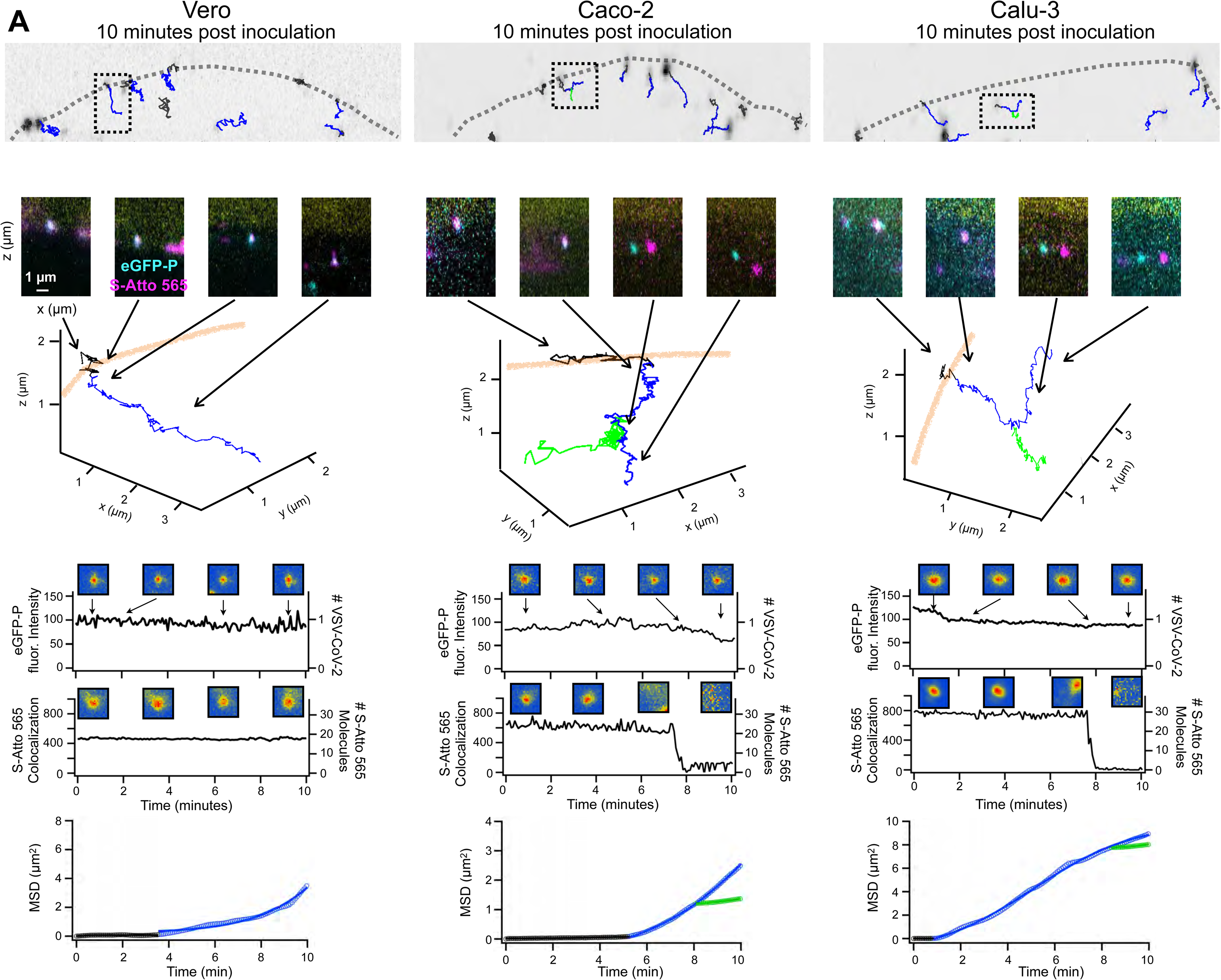
VSV-eGFP-P-SARS-CoV-2 uptake in Vero E6, Caco-2 and Calu-3 cells. Trajectories overlayed onto a 2 µm projection though the x-axis of a cell imaged using LLSM microscopy with whole cell volumes collected every 4.7 seconds. Over the course of 10 minutes virus was observed to be on the cell surface (black trajectories), internalize into the cell with both eGFP-P and Atto 565 dye co-localizing (blue trajectories), with some eGFP-P observed to separate from Atto 565 signal (green). Example images of boxed trajectories shown below with complete orthogonal projections underneath. The intensity of the eGFP-P and the intensity of Atto 565 co-localizing with the eGFP-P signal are showing as well as the MSD analysis of the trajectory calculated localize when the particles are co-localized to the surface, inside the cell following the trajectory of the Atto 565 signal or following the trajectory of the eGFP-P signal after loss of co-localization with Atto 565.

**Figure S4.**
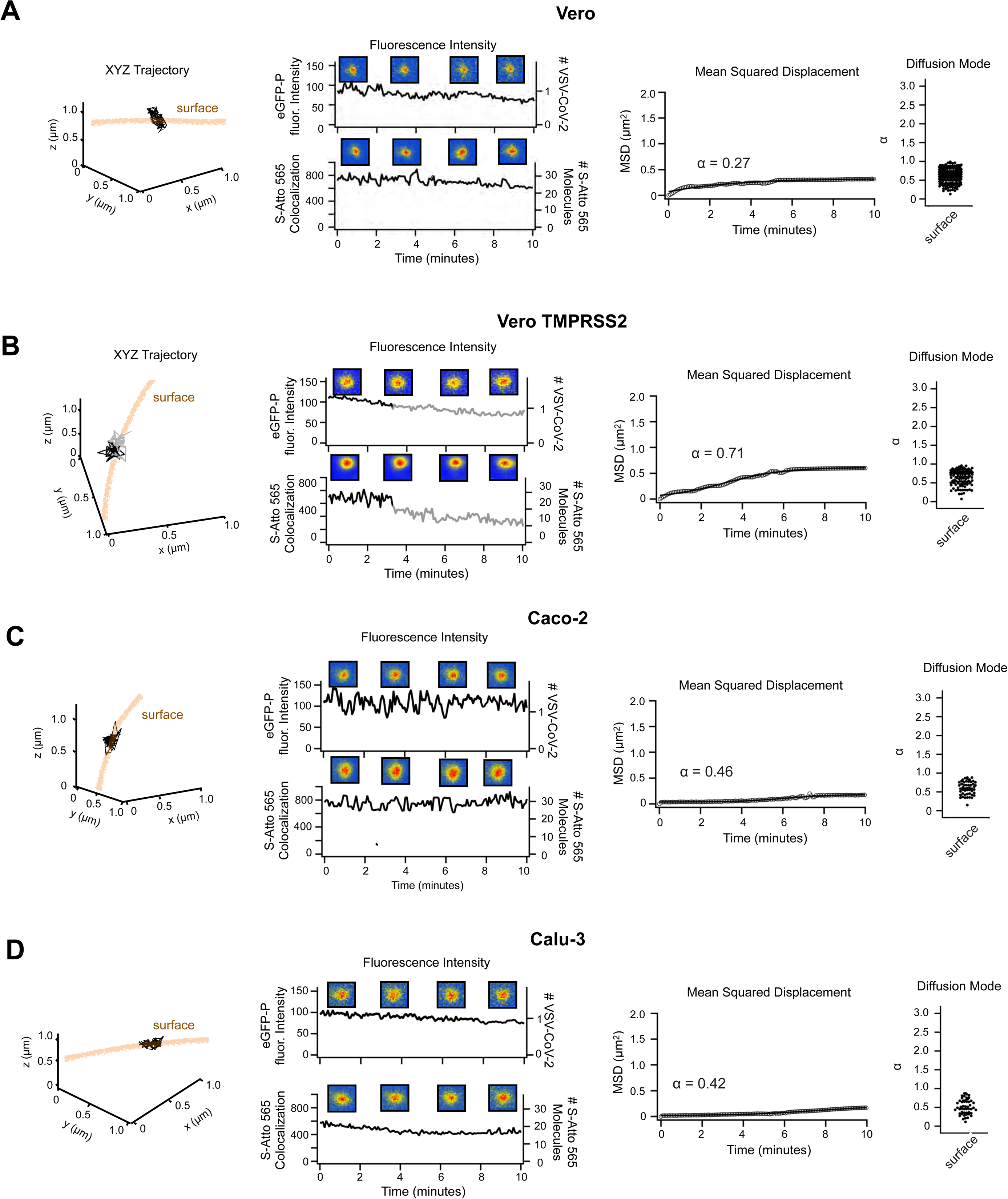
VSV-eGFP-P-SARS-CoV-2 surface colocalized trajectories. Example events of VSV-eGFP-P-SARS-CoV-2-Atto-565 co-localized to the cell surface from Vero E6 **(A)**, Vero E6 TM **(B)** Caco-2 (**C)** and Calu-3 **(D)** cells. A complete orthogonal projection (left), the intensity of eGFP-P and co-localization with Atto 565 spots (middle left), the MSD analysis (middle right), and the alpha values of every virus trajectory during their colocalization at the cell surface in the respective cell type.

**Figure S5.**
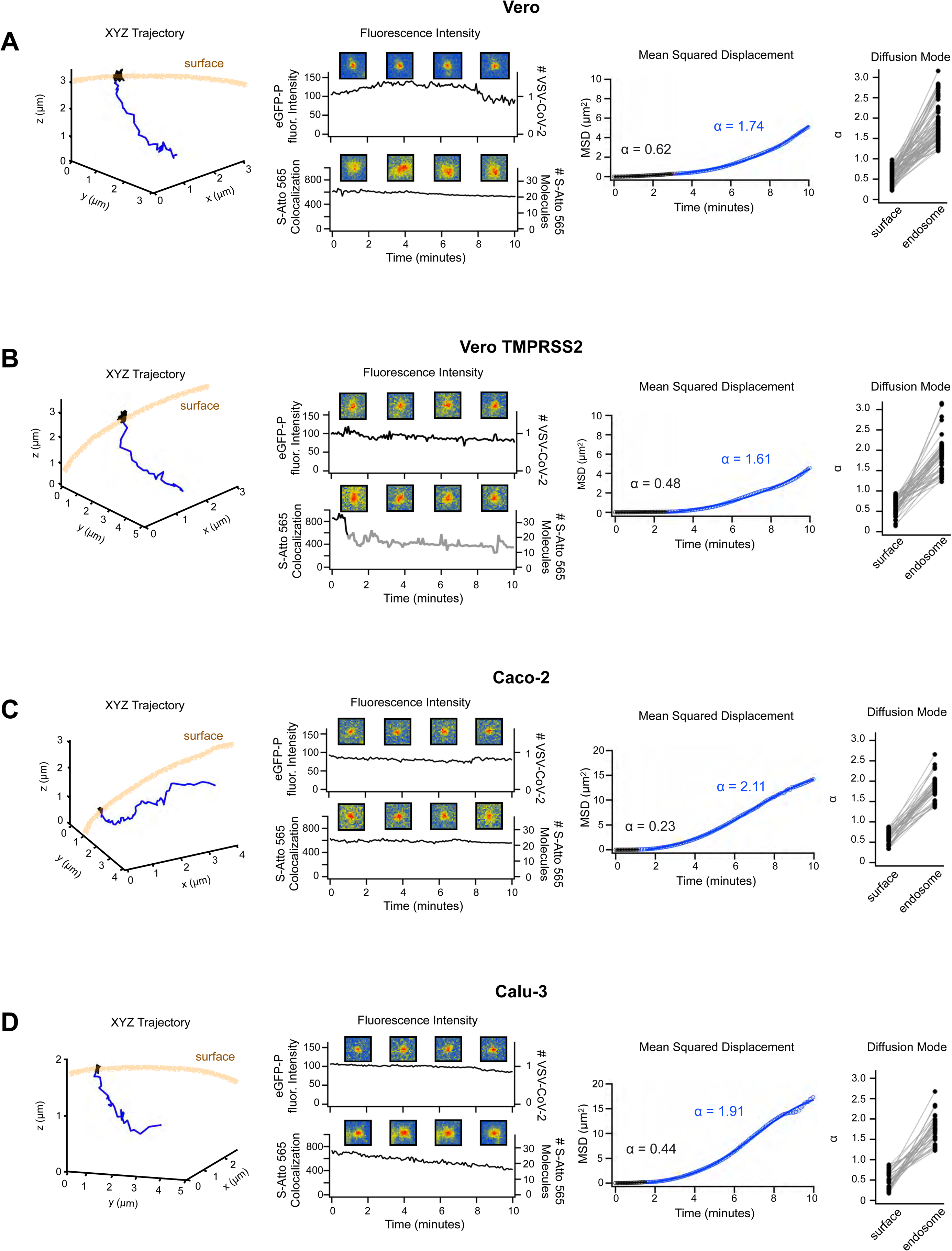
VSV-eGFP-P-SARS-CoV-2 endocytosis trajectories. Example events of VSV-eGFP-P-SARS-CoV-2-Atto-565 co-localized to the cell surface and subsequent endocytosis into Vero E6 **(A)**, Vero TMPRSS2, **(B)** Caco-2 **(C)** and Calu-3 **(D)** cells. A complete orthogonal projection (left), the intensity of eGFP-P and co-localization with S Atto 565 spots (middle left), the MSD analysis (middle right), and the alpha values (right) of every trajectory when it was co-localized to the cell surface then to the alpha value after it entered the cell volume for every virus with this behavior in the respective cell type.

**Figure S6.**
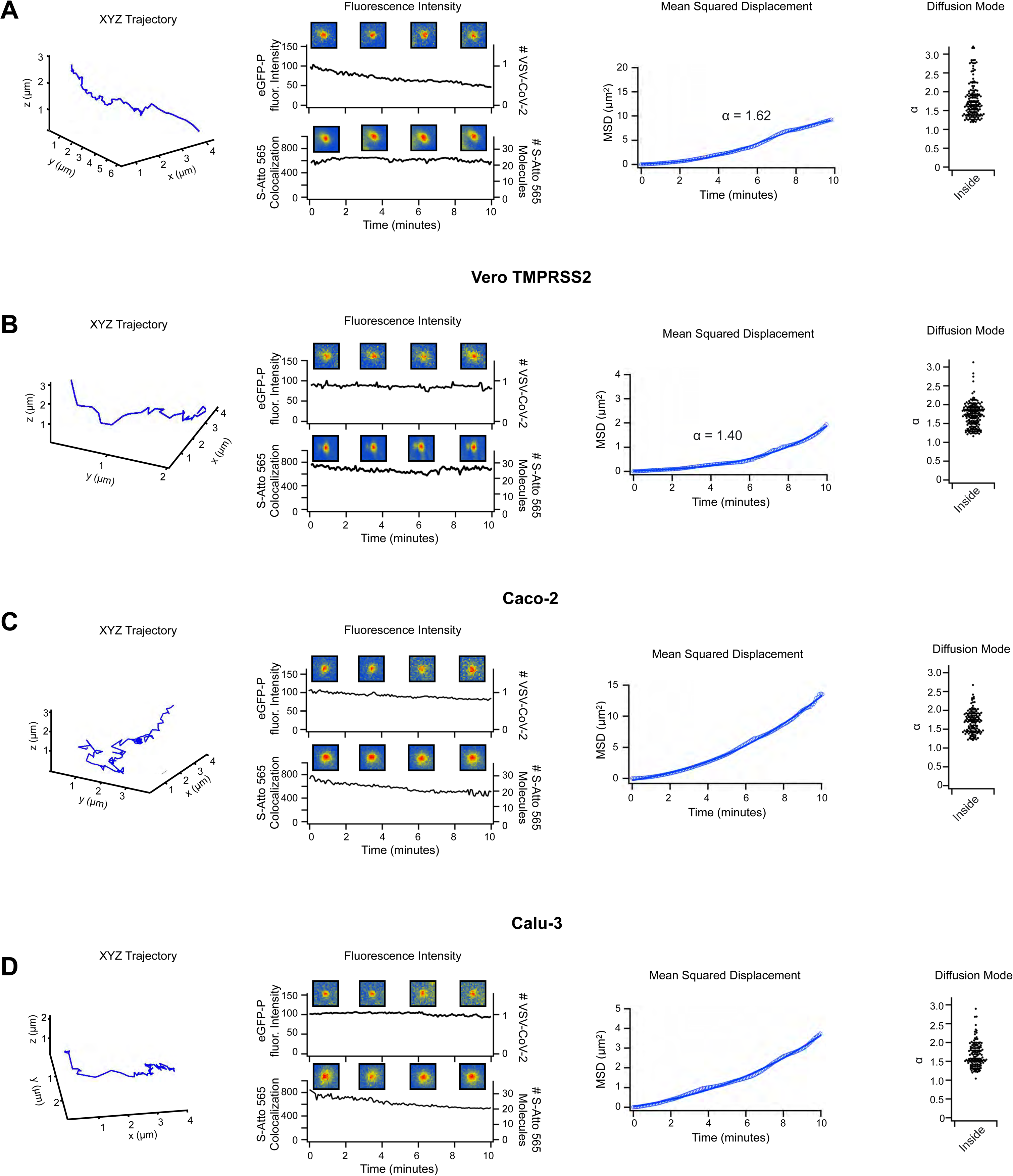
VSV-eGFP-P-SARS-CoV-2 inside endosomes. Example events of VSV-eGFP-P-SARS-CoV-2-Atto-565 co-localized with eGFP-P and Atto565 colocalized inside Vero E6 (A), Vero TMPRSS2 (B) Caco-2 (C) and Calu-3 (D) cells. A complete orthogonal projection (left), the intensity of eGFP-P and co-localization with S Atto 565 spots (middle left), the MSD analysis (middle right), and the alpha values (right) of every trajectory exemplifying this behavior in the respective cell type.

**Figure S7.**
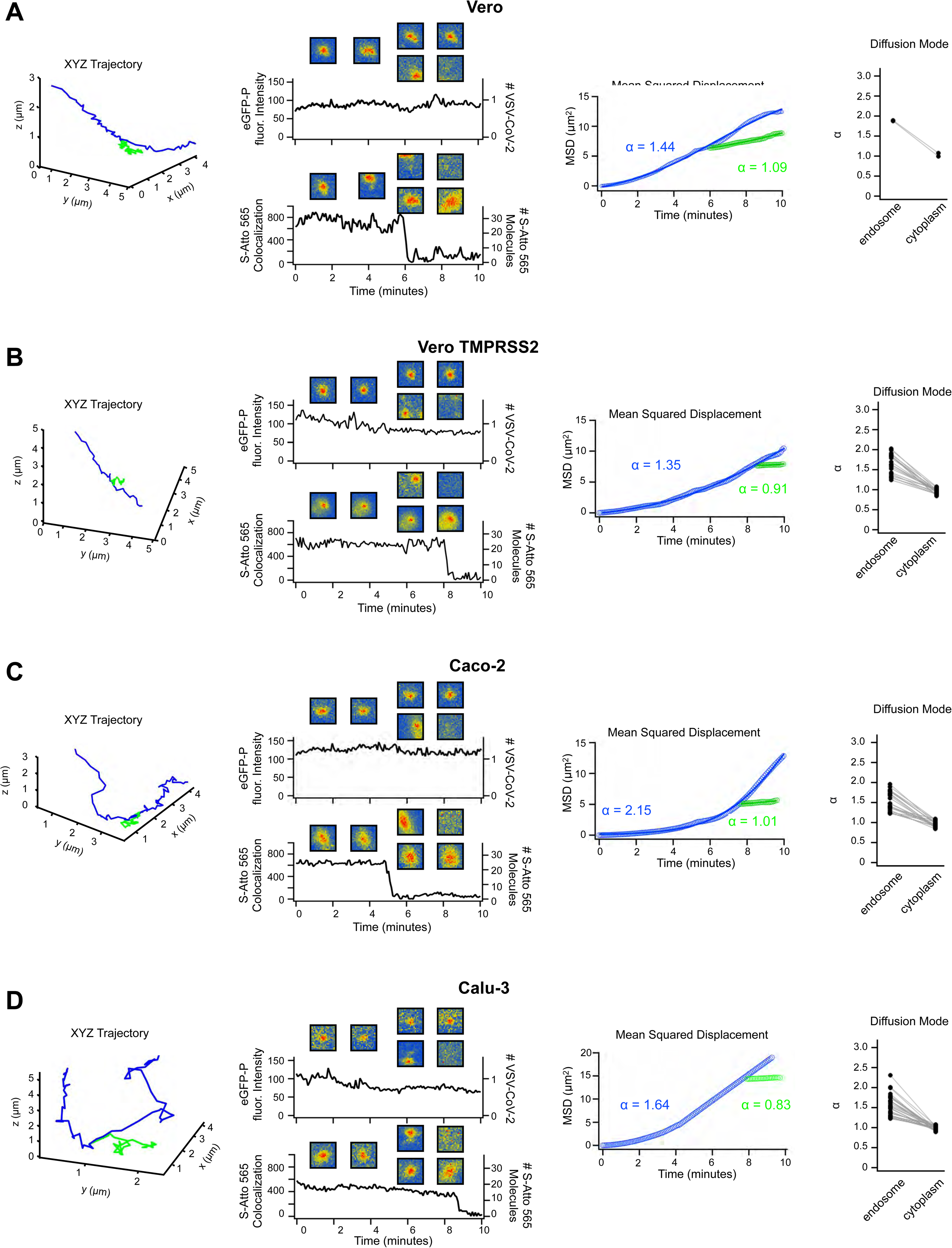
VSV-eGFP-P-SARS-CoV-2 fusion events from endosomes. Example events of VSV-eGFP-P-SARS-CoV-2-Atto-565 inside the cell with initial colocalization and subsequent separation of the eGFP-P and Atto 565 signals from Vero E6 **(A)**, Vero TMPRSS2, **(B)** Caco-2, **(C)** and Calu-3 **(D)** cells. A complete orthogonal projection (left), the intensity of eGFP-P and co-localization with Atto 565 spots (middle left), the MSD analysis (middle right), and the alpha values (right) of every trajectory exemplifying this behavior in the respective cell type.

**Figure S8.**
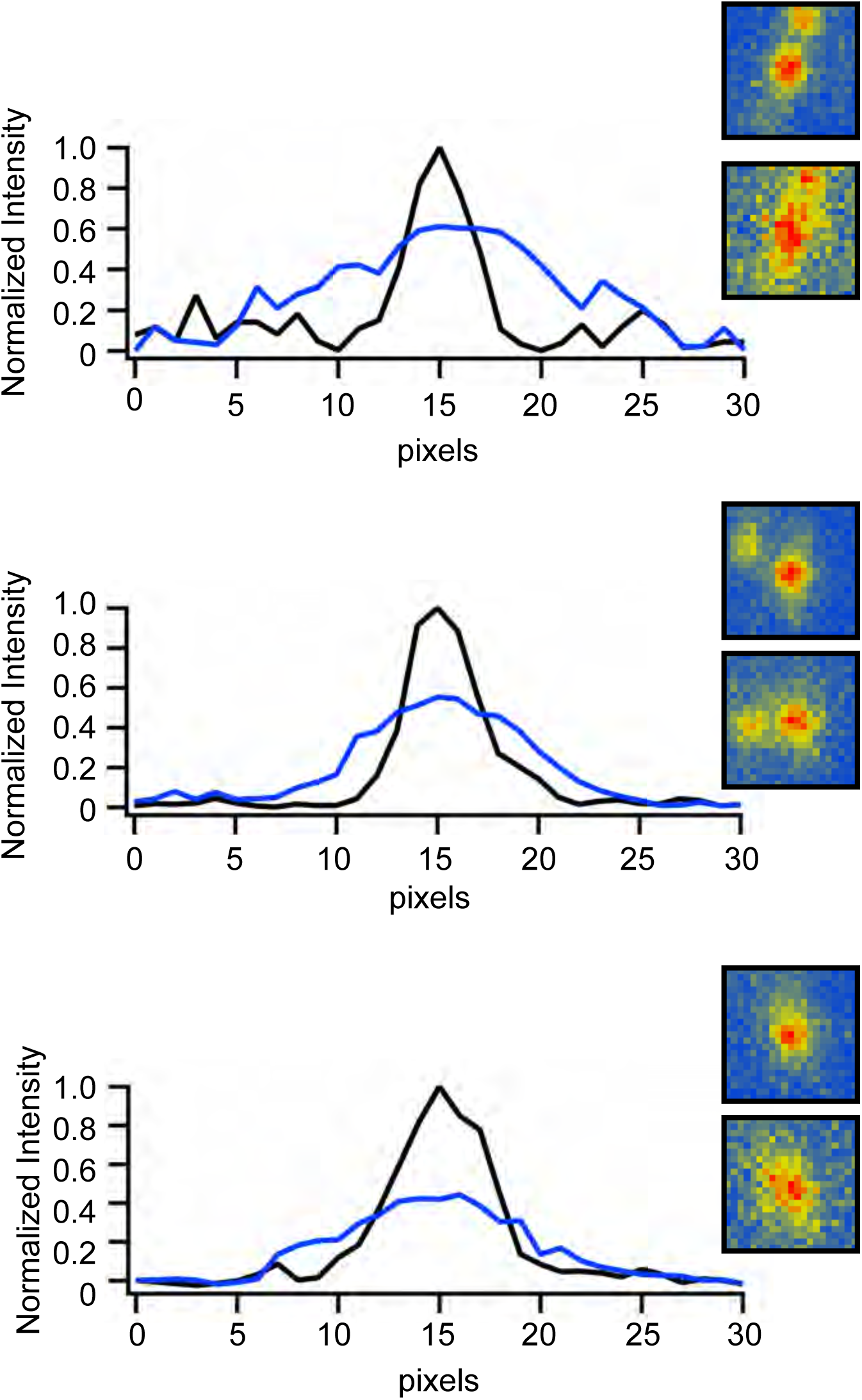
The spread of the S associated Atto 565 signal after fusion between the viral envelope and endosomal membrane. Cross section showing three representative examples of the intensity distribution of the S associated Atto 565 signal from VSV-eGFP-P-SARS-CoV-2-Atto 565 before (black) and after (blue) separation of eGFP-P. Insets show images of the events used for the intensity profiles.

**Figure S9.**
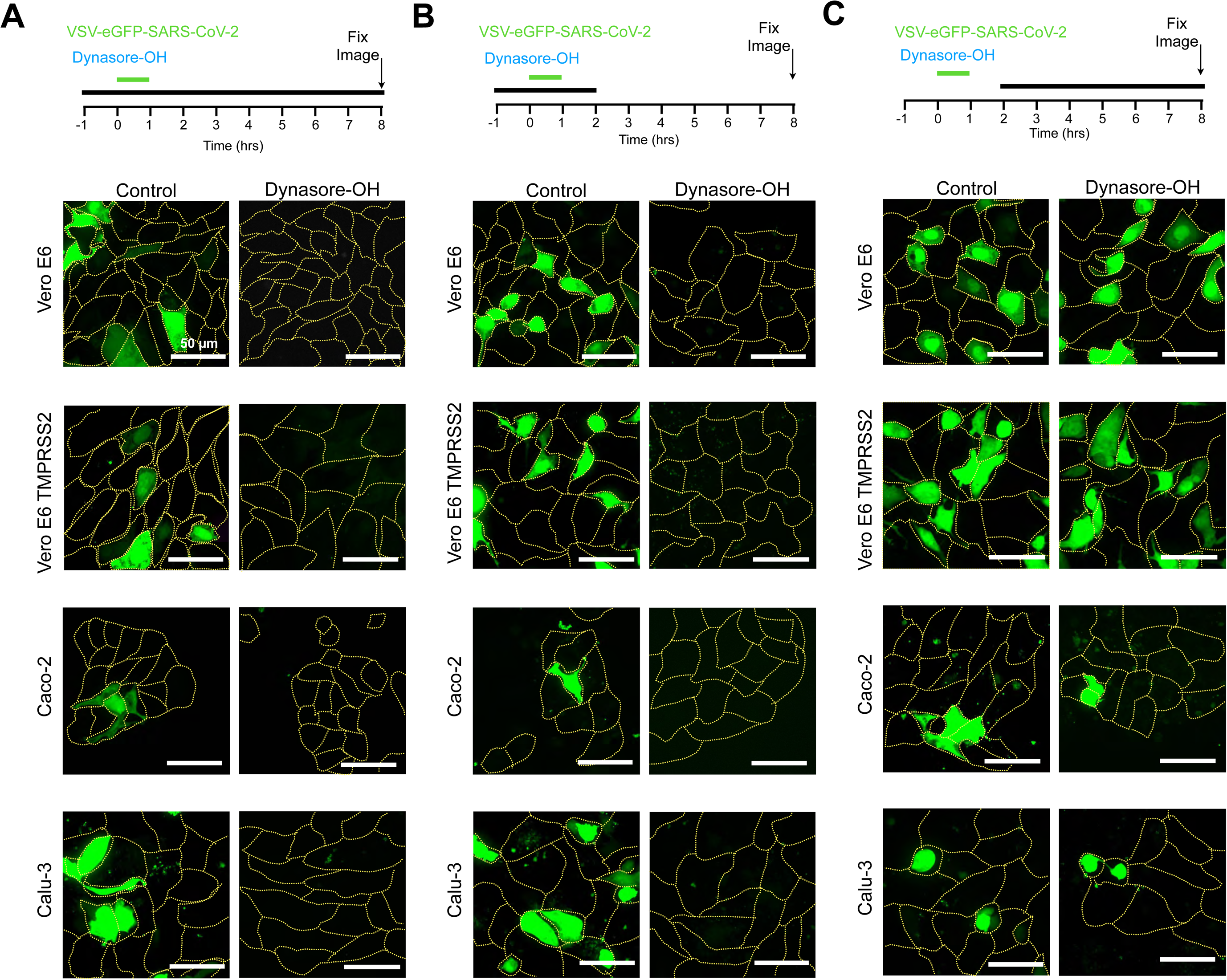
Chemical inhibition of dynamin inhibits an early step of VSV-SARS-CoV-2 infection. Dynasore at 40 uM was incubated with the respective cell type for **(A)** beginning 1 hour prior to addition of VSV-SARS-CoV-2 throughout the experiment, **(B)** beginning 1 hour prior to addition of VSV-SARS-CoV-2 until 1 hour after virus was removed, and **(C)** beginning 1 hour after the washing away of unbound VSV-SARS-CoV-2.

**Figure S10.**
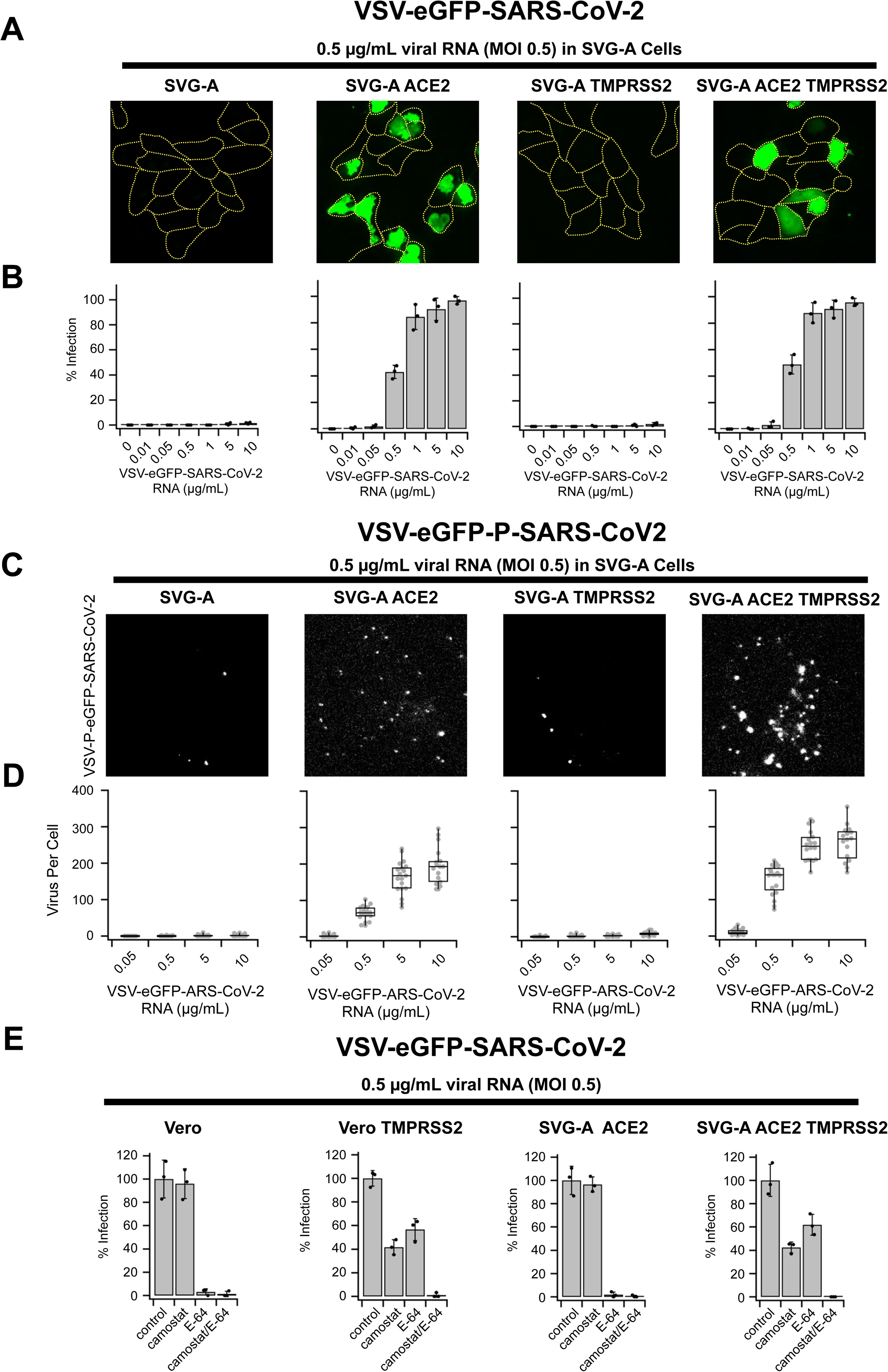
VSV-eGFP-SARS-CoV-2 infection of SVG-A cells endogenously expressing EEA1-Scarlett and NPC1-Halo. **(A)** Max intensity projections of example images taken with 1 µm spacing for 20 µm. **(B)** Quantification of VSV-eGFP-SARS-CoV-2 infection in parental (right), ACE2 expressing (middle right), TMPRSS2 expressing (middle left), and ACE2 combined with TMPRSS2 expressing (left) SVG-A cells. **(C)** Max intensity projection of 10 planes taken with 0.27 µm spacing of VSV-eGFP-P-SARS-CoV-2 uptake and quantification **(D)** Whole cell volumes were taken with each data point representing number of virus per cell for parental (right), ACE2 expressing (middle right), TMPRSS2 expressing (middle left), and ACE2 combined with TMPRSS2 expressing (left) SVG-A cells.

**Figure S11.**
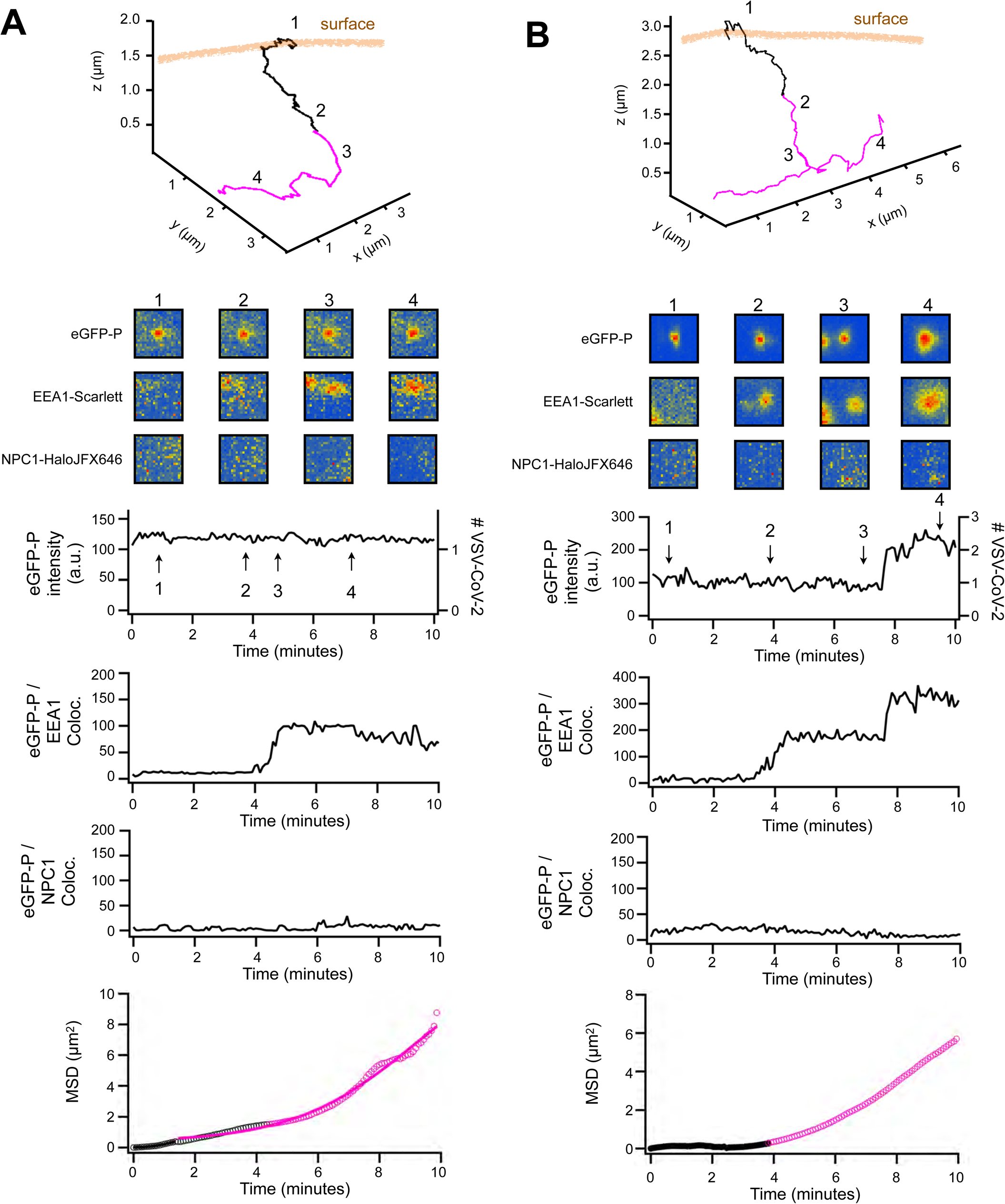
Endocytosis and arrival of VSV-eGFP-P-SARS-CoV-2 to early endosomes. **(A, B)** Example events with orthogonal projections (top), insets centered around the trajectory coordinates at discrete time points, eGFP-P fluorescence, and co-localization intensities for the EEA1-Scarlett and NPC1-HaloJFX646 channels, and the MSD analysis of the event. **(A)** Depicts endocytic uptake of virus and accumulation in an EEA1 positive compartment over ∼2 minutes after internalization. **(B)** Depicts virus internalization and accumulation in an EEA1 positive compartment with subsequent fusion with another EEA1-Scarlett positive endosome.

**Figure S12.**
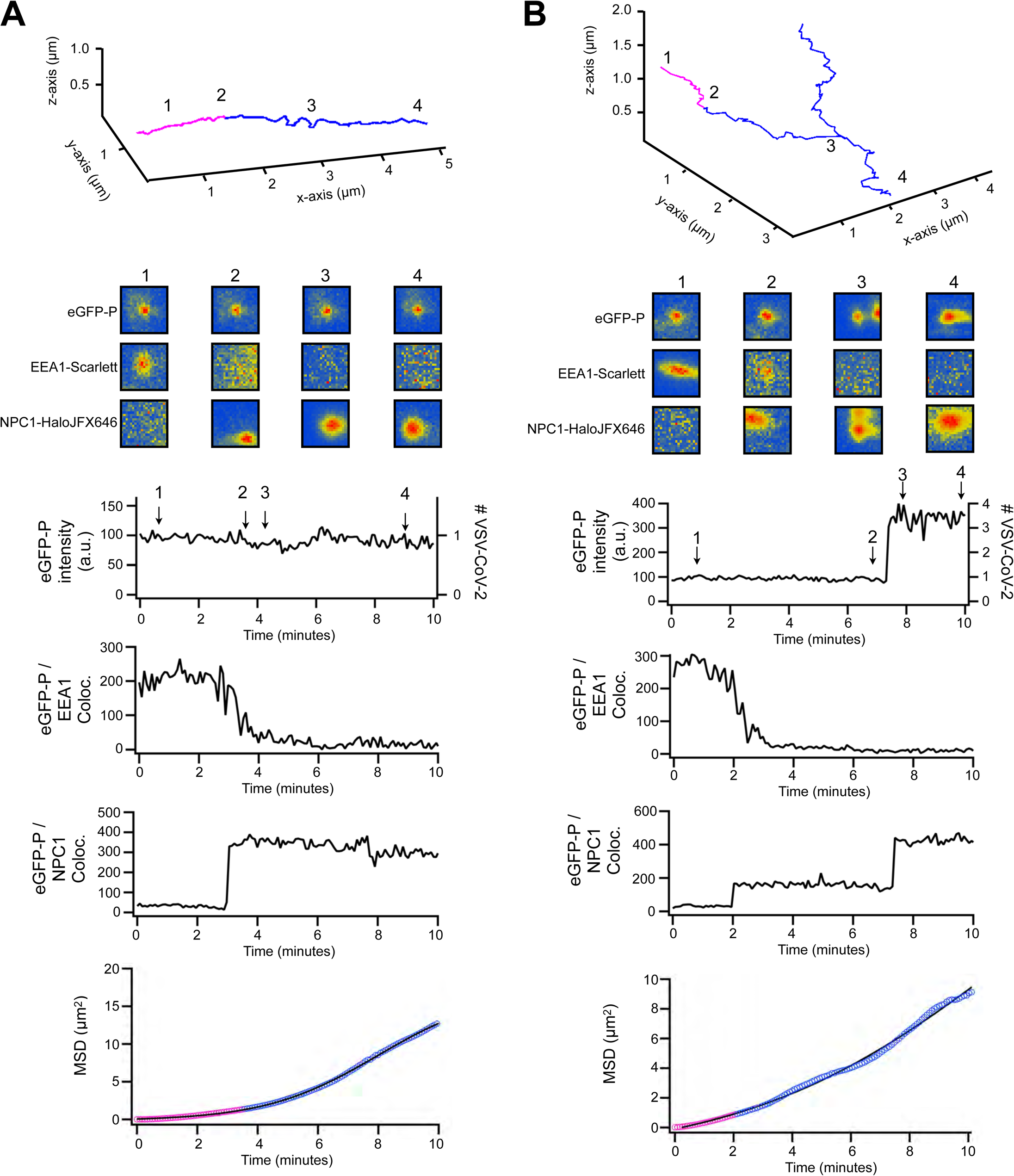
Trafficking of VSV-eGFP-P-SARS-CoV-2 from early to late endosomes. **(A, B)** Example events with orthogonal projections (top), insets centered around the trajectory coordinates at discrete time points, eGFP-P fluorescence, and co-localization intensities for the EEA1-Scarlett and NPC1-HaloJFX646 channels, and the MSD analysis of the event. **(A)** An event depicting virus in an early endosome with subsequent loss of EEA1-Scarlett over a 1-minute time period and the rapid increase in NPC1-HaloJFX646 indicative of a fusion event with a late endosome. (B) A virus particle in an early endosome showing loss of EEA1 concomitant with merging with a late endosomal, NPC1-HaloJFX646 positive compartment. This particle then diffuses and undergoes a subsequent fusion event with a NPC1-HaloJFX646 positive endosome that contains eGFP-P corresponding to ∼2 viruses.

**Figure S13.**
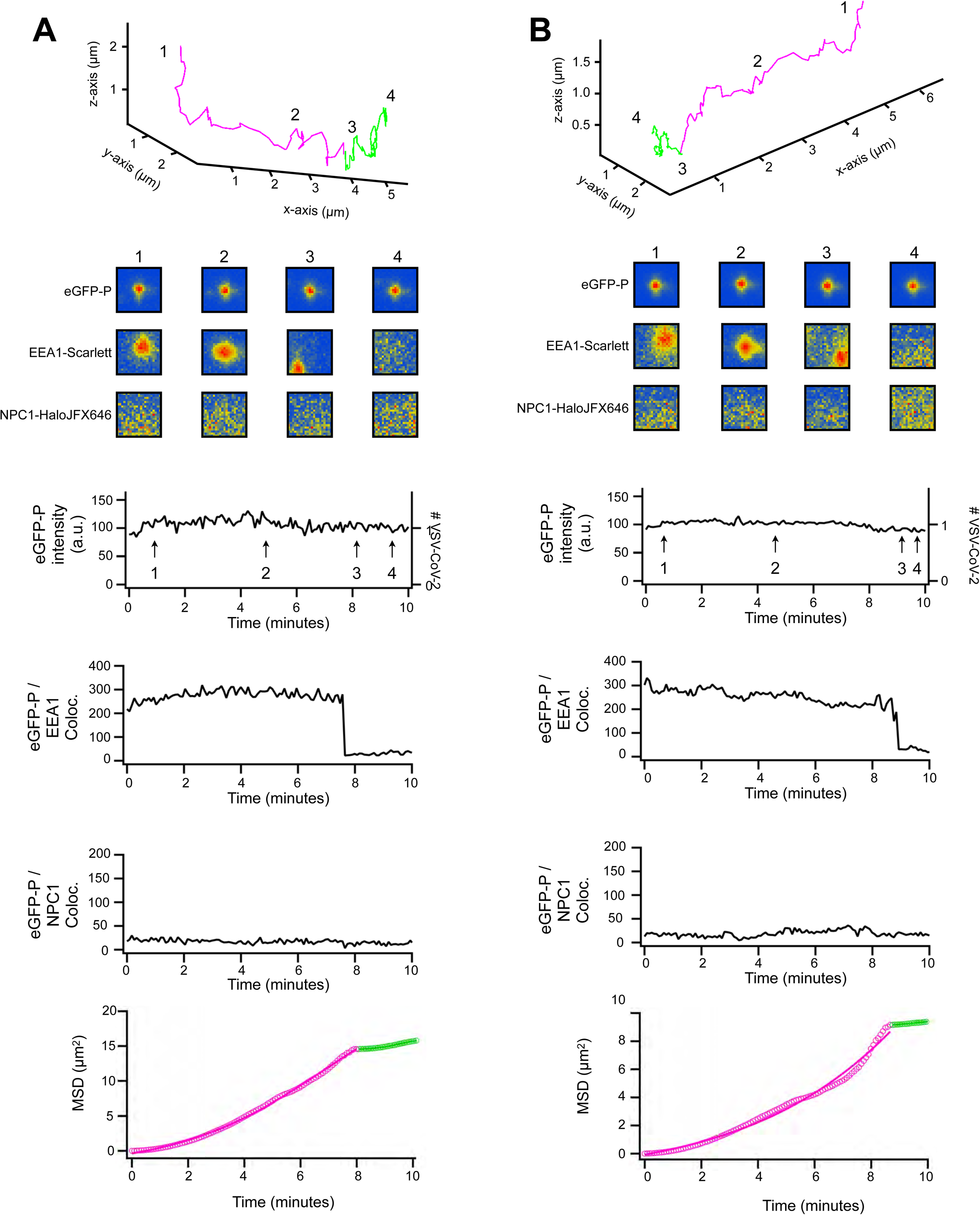
Release of VSV-EGFP-P-SARS-CoV-2 from an early endosome. **(A, B)** Example events with orthogonal projections (top), insets centered around the trajectory coordinates at discrete time points, eGFP-P fluorescence, and co-localization intensities for the EEA1-Scarlet and NPC1-HaloJFX646 channels, and the MSD analysis of the event. Both examples show a single virus in EEA1-Scarlett positive endosome.

**Figure S14.**
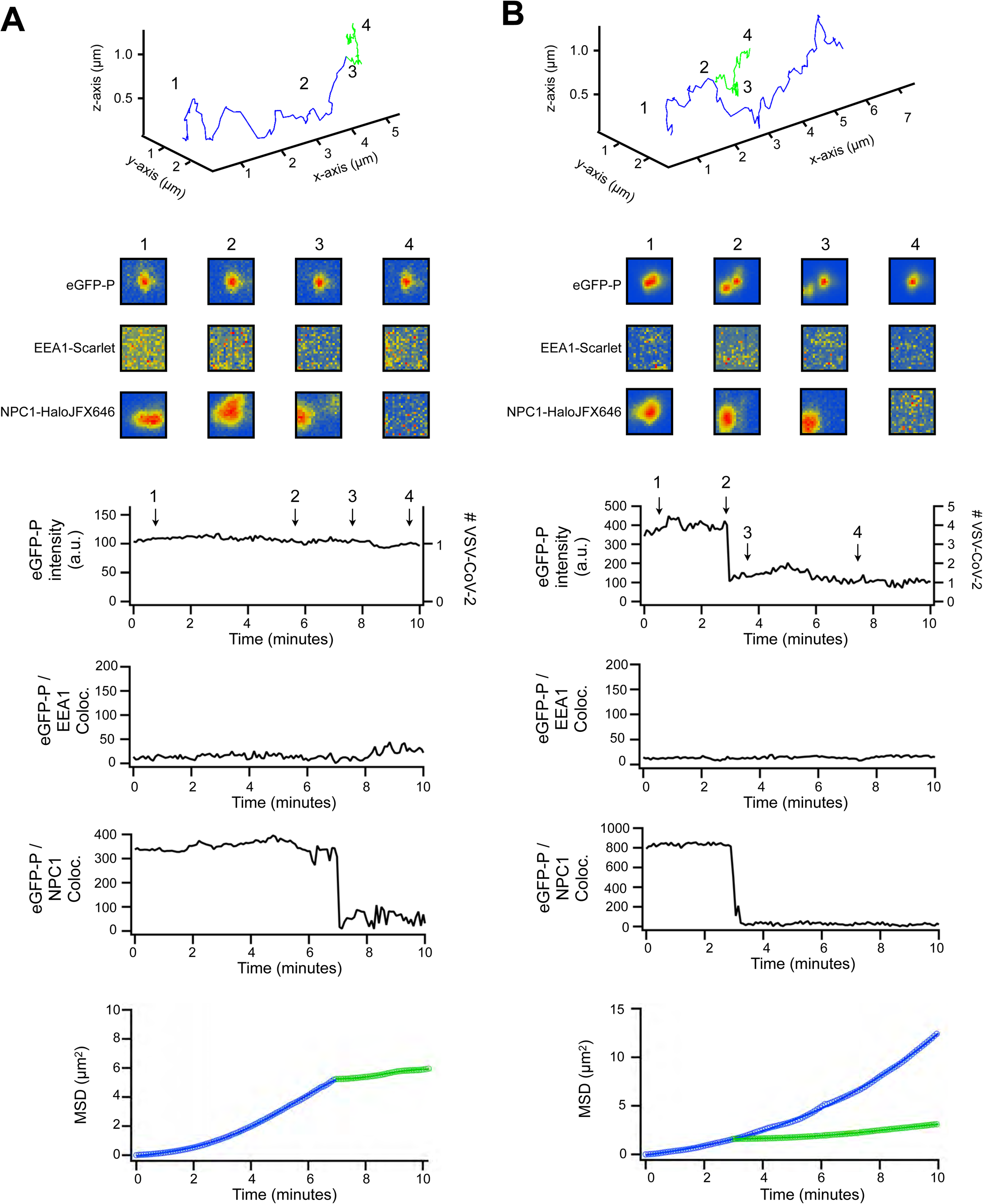
Release of VSV-eGFP-P-SARS-CoV-2 from a late endosome. **(A, B)** Example events with orthogonal projections (top), insets centered around the trajectory coordinates at discrete time points, eGFP-P fluorescence, and co-localization intensities for the EEA1-Scarlett and NPC1-HaloJFX646 channels, and the MSD analysis of the event. **(A)** Shows a release event of a single particle from an NPC1-HaloJFX646 positive late endosome. **(B)** Shows a NPC1-HaloJFX646 positive late endosome with fluorescence corresponding to roughly 4 viruses. The signal corresponding to 1 virus releases from the endosome and exhibits Brownian motion while the signal from the remainder continues to traffic with the late endosome.

**Figure S15:**
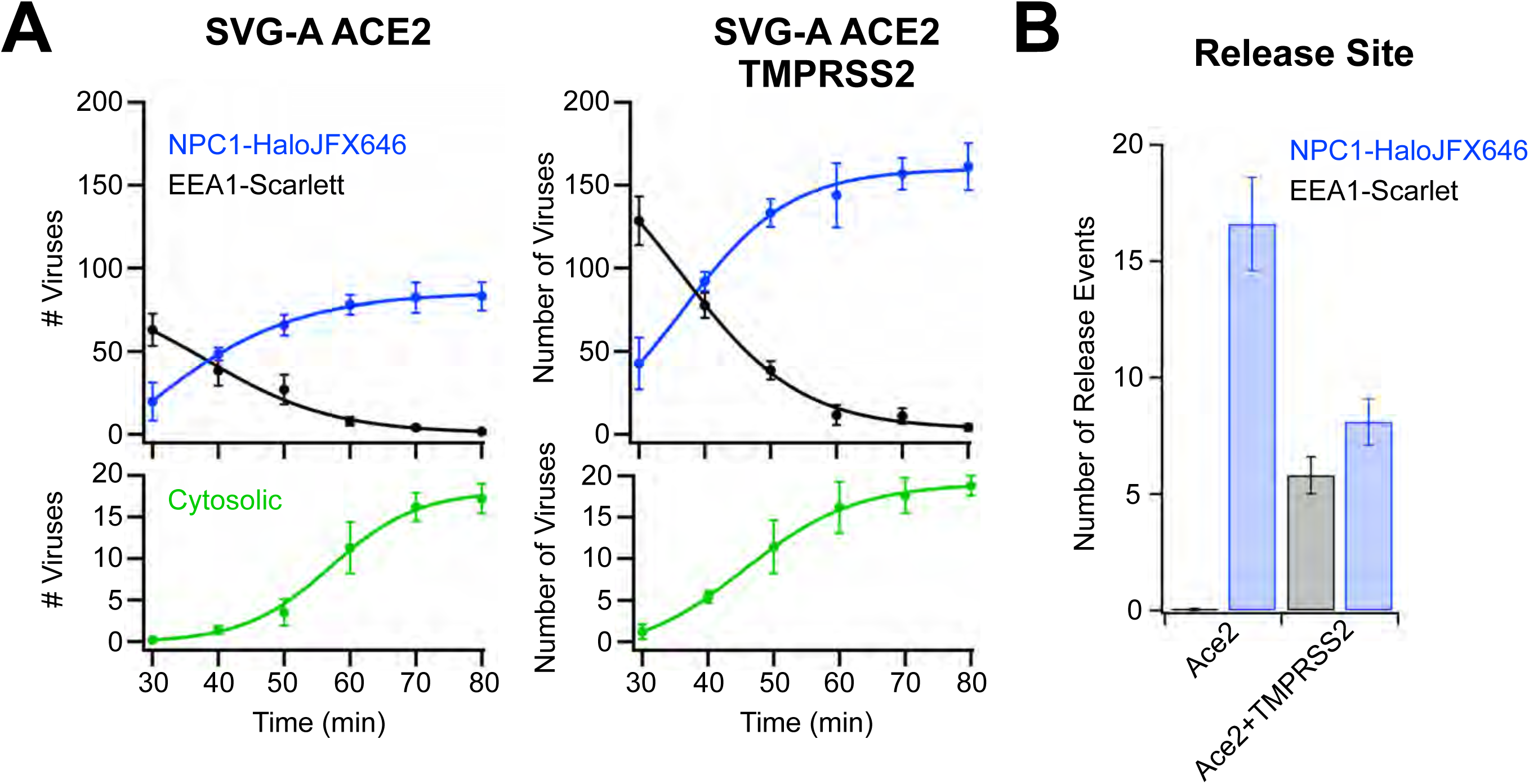
Analysis of the location of VSV-eGFP-P-SARS-CoV-2 throughout the endosomal pathway. **(A)** Discrete timepoint in 10-minute intervals of the location of VSV-eGFP-P-SARS-CoV-2 in gene-edited SVG-A EEA1-Scarlett NPC1-HaloJFX646 cells imaged using LLSM from the data in Fig 2. The average localization of the virus in either early or late endosomes, and the fraction of viral contents released into the cytosol is shown for cells expressing ACE2 (left) or ACE2 and TMPRSS2 (right). Each point is an average of 5 cells at that timepoint with errors being S.E.M. **(B)** All observed release events from a single coverslip spanning 5 cells imaged consecutively were pooled to examine the number of observed fusion events in EEA1 positive early endosome or NPC1 positive late endosomes/lysosomes. Each condition is an average of 5 coverslips where 5 cells were imaged consecutively for 10 minutes from 30 to 80 minutes post inoculation. Error bars are S.E.M.

**Figure S16:**
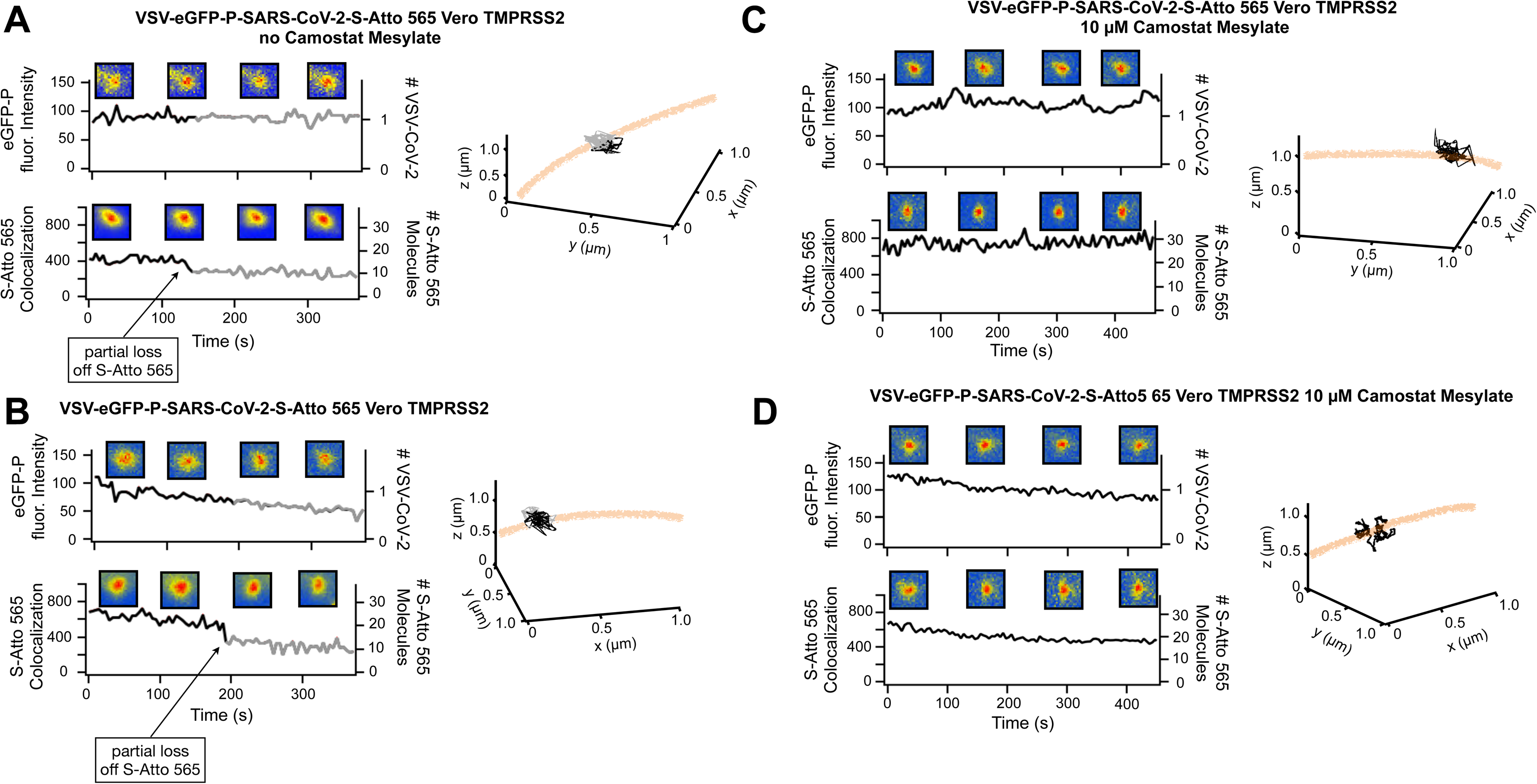
Loss of S-tagged Atto565 is TMPRSS2 dependent. **(A, B)** Intensity profile and orthogonal projections of single VSV-eGFP-P-SARS-CoV-2-Atto 565 on the surface of Vero TMPRSS2 cells showing a characteristic stepwise loss of Atto 565 signal. **(C, D)** As in A, B except that cells were pretreated for 1h with 10 µM camostat mesylate, and particles showed no stepwise loss in Atto 565 signal.

**Figure S17.**
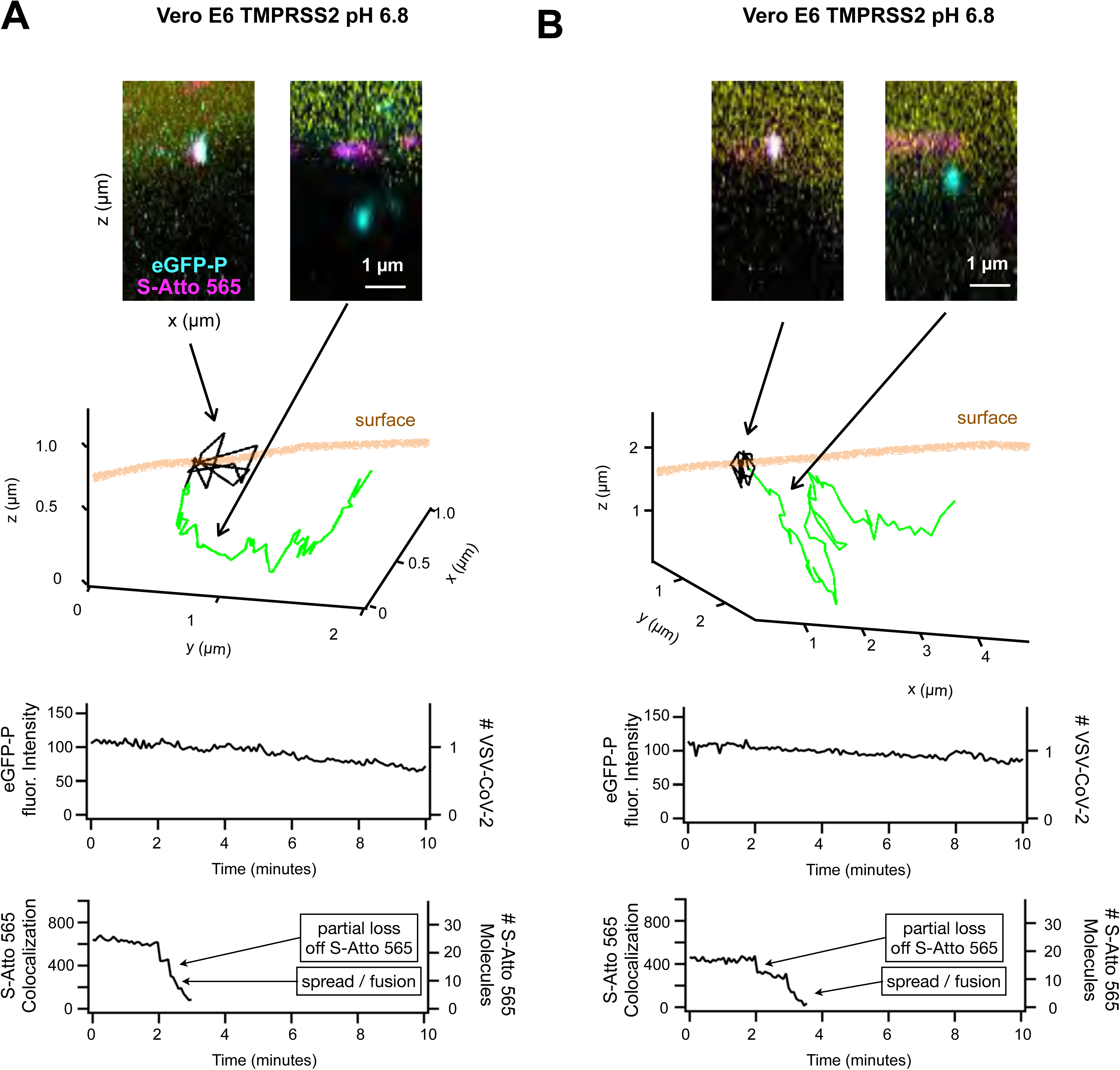
TMPRSS2 dependent surface fusion in the presence of slightly acidic extracellular media. **(A, B)** Orthogonal projections (top) show location of VSV-eGFP-P-SARS-CoV-2-S-Atto 565 on the cell surface (black) where eGFP-P is released into the cytosol (green). Example images of a particle before and after the fusion event evident by separation of the eGFP (light blue) from the Atto565 (magenta) on the surface with subsequent spread of the Atto565 signal on the plasma membrane of a Vero TMPRSS2 cell at extracellular pH 6.8. Intensity of the events over time are shown where the spread and decay of the S Atto 565 signal occurs following the stepwise drop indicating release of S1 from the S2 of the spike protein.

**Figure S18.**
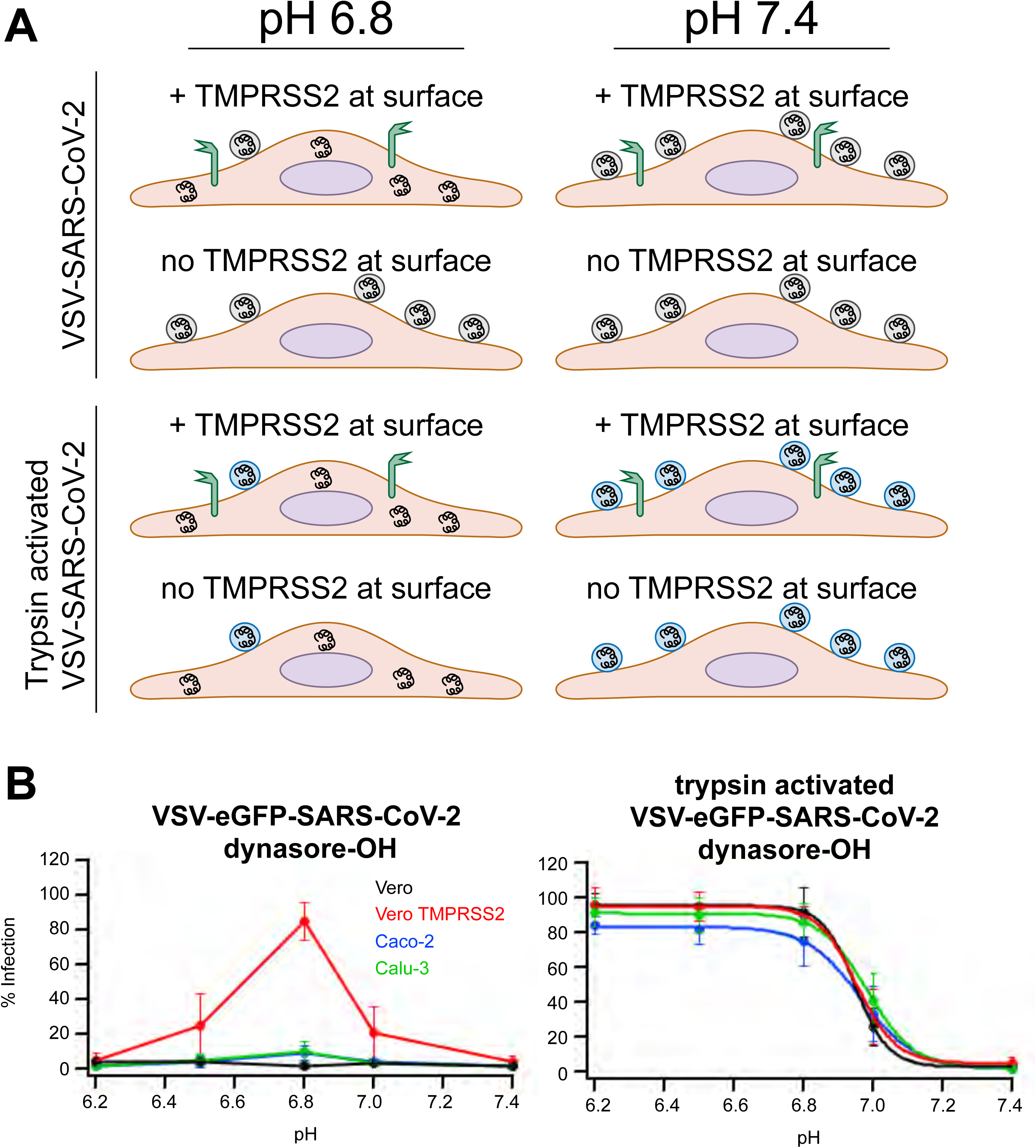
Proteolytic activation of S and the pH dependence of VSV-eGFP-SARS-CoV-2 infection in different cell types. **(A)** Schematic representation of a pH bypass experiment comparing the behavior of VSV-eGFP-SARS-CoV-2 with or without trypsin activation. **(B)** Vero, Vero TMPRSS2, Caco-2 and Calu-3 cells were treated with 40 uM dynasore-OH for 1 hr prior to exposure to VSV-eGFP-SARS-CoV-2 at an MOI of ∼ 0.5 in media of varying pH. The relative level of infection is shown compared to infection in the absence of dynasore-OH (left graph). Pretreatment of virus with trypsin (right graph) rescues infection in all cell types in the presence of dynasore-OH.

**Figure S19.**
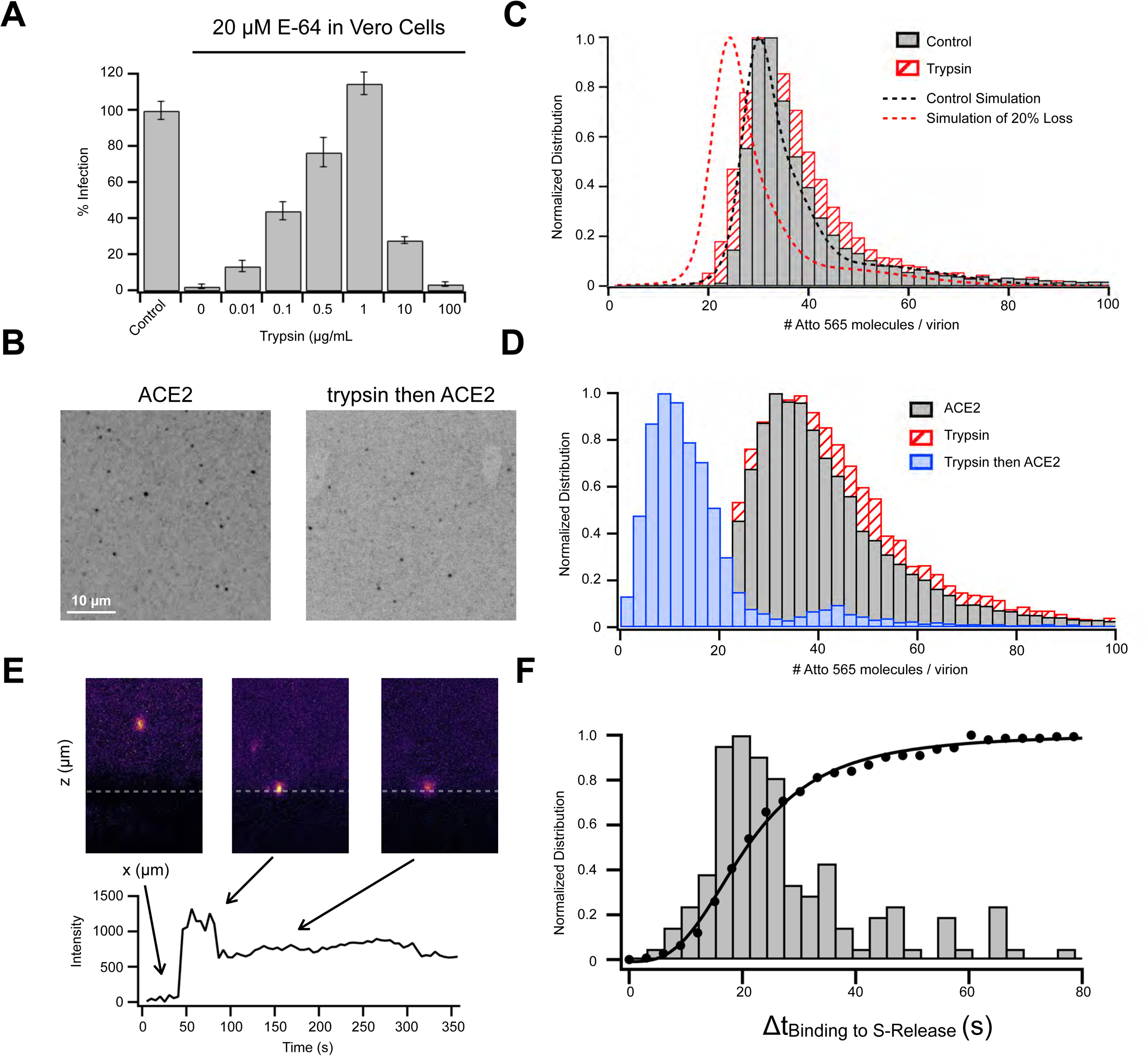
Reconstitution of S1 release from VSV-eGFP-SARS-CoV-2-Atto 565 particles. **(A)** VSV-eGFP-SARS-CoV-2-Atto 565 infects Vero cells in the presence of the host cell cathepsin protease inhibitor (20 µM E-64) when viral particles are pretreated with trypsin for 30 min at 37°C (followed by inhibiting trypsin with Aprotinin). **(B-D)** Calibration of the fluorescence intensity of single Atto 565 molecules using photobleaching and then imaging thousands of VSV-eGFP-SARS-CoV-2-Atto 565 that were allowed to adsorb on poly-lysine coated coverslips showed that trypsin cleaved virus needs to bind ACE2 to release the S1 fragment. **(B)** Shows example images of a max intensity projection of 10 planes spaced every 0.27 µm obtained by spinning-disc confocal microscopy of virus treated with ACE2 alone (left) or first treated with trypsin and then incubated with ACE2. **(C)** Number of Atto 565 per virus of an experiment with and without trypsin cleavage. A numeric simulation was run to determine the distribution of a data set with similar labeling with and without a 20% loss in signal to mimic what was observed in for Atto 565 signal loss in the LLSM experiments (Figure 3, S16, S17). **(D)** Fluorescence intensity distribution of Atto 565 signal shows that the addition of ACE2 only causes a loss of Atto 565 signal if the virus was pretreated with trypsin. All histograms are single experiments with distributions representing more than 8,000 per experiment. Each experiment was done in triplicate with the peak intensity of distributions determined to be 38.2 ± 3.4, 39.7 ± 4.1, 38.5 ± 1.6, and 9.2 ± 2.4 Atto 565 molecules per virus for control, trypsin, ACE2, or trypsin followed by ACE2 treatments respectively. **(E)** Trypsin cleaved VSV-SARS-CoV-2-Atto 565 was rapidly added to Vero cells (lacking any surface protease) using a microfluidic flow cell and after binding a stepwise drop corresponding to ∼25% signal was observed after a delay (F) which was measured for 130 virus binding events where most step drops occurred between 8-30 seconds after virus binding.

**Figure S20.**
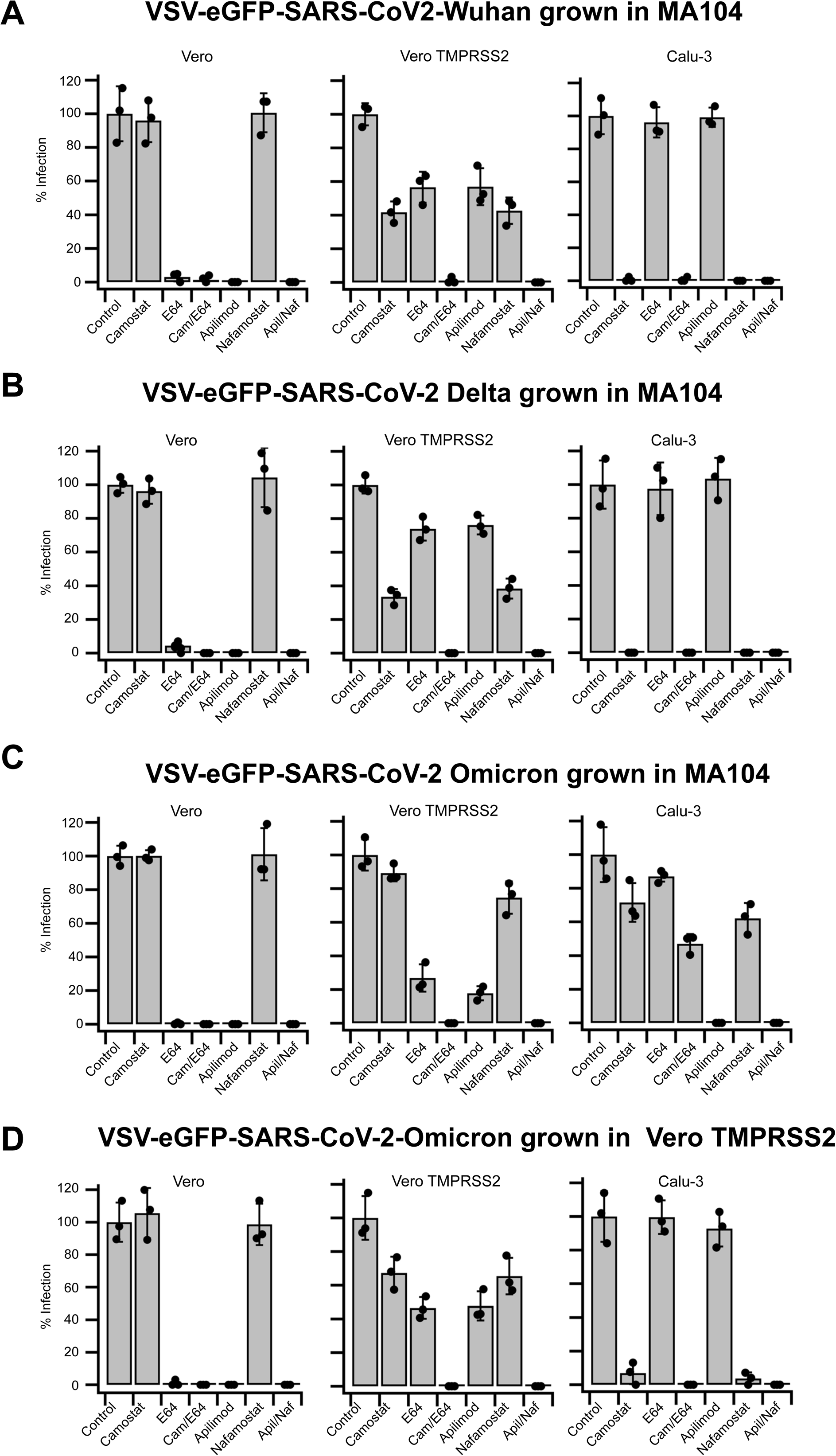
Drug sensitivities of VSV-SARS-CoV-2 spike variants show that furin cleavage determine proteolytic entry route. **(A-D)** VSV-eGFP-SARS-CoV-2 infection of spike variants including Wuhan **(A)**, Delta **(B)**, and Omicron **(C, D)** with different levels of furin cleavage where virus grown in MA104 cells **(C)** had defective furin cleavage while virus grown in Vero TMPRSS2 cells **(D)** were where furin cleaved (Fig. 4). Cells were treated with indicated drugs for 1 hour, infected with virus for 1 hr, and then washed while maintaining the concentration of the drug constant. After 8 hours cells were fixed, stained with WGA-Alexa647, and imaged for eGFP expression using a spinning-disc confocal microscope. The sensitive to TMPRSS2 was determined by treatments with 10 µM camostat mesylate or 100 nM nafamostat mesylate. The sensitivity to cathepsins was determined with 20 µM E-64 while the endosomal trafficking route necessary for cathepsin dependent entry was targeted with 100 nM of the inhibitor for PIKfyve, apilimod. Combinations of 10 µM camostat mesylate and 20 µM E-64 or 100 nM nafamostat mesylate and 100 nM apilimod completely abolished infection in all cell types as previously described (*2, 3*). Cells were infected at an MOI of ∼ 0.5 which was 0.5 µg/mL virus RNA for all spike variants in Vero and Vero TMPRSS2 and 5 µg/mL in Calu-3 cells, except for the furin defective VSV-eGFP-SARS-CoV-2-Omicron **(C)** which required 30 µg/mL viral RNA to achieve and MOI of 0.5 in Calu-3 cells.

**Figure S21.**
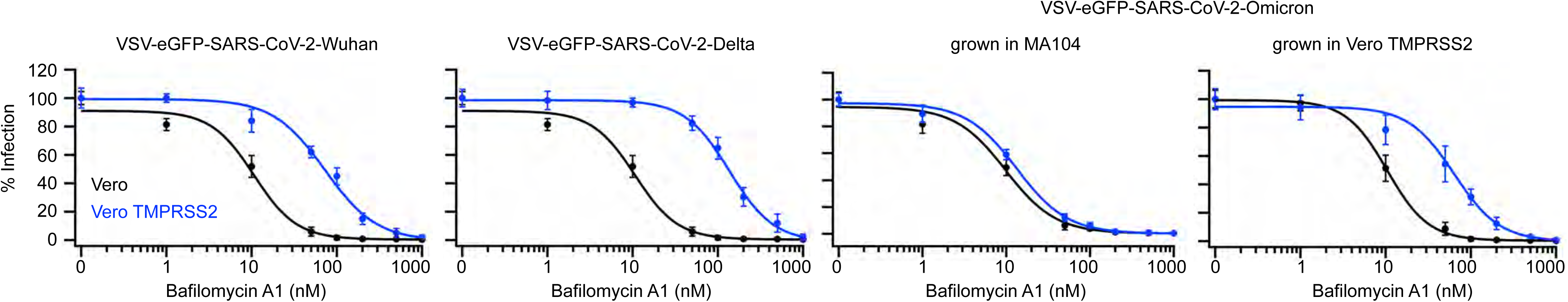
Inhibition of endosomal acidification blocks VSV-SARS-CoV-2 infection for all variants with a shift to a lower affinity in the presence of TMPRSS2 protease. Cells where incubated with indicated concentration of bafilomycin A1 for 1 hour, indicated VSV-eGFP-SARS-CoV-2 spike variant was added for 1 hour, unbound virus was then washed away, and 7 hrs later cells were labeled with WGA-Alexa647, fixed, and imaged for eGFP expression using a spinning-disc confocal. Bafilomycin A1 was kept constant throughout the time course of the infection assay.

## MATERIAL AND METHODS

### Materials

The following reagents were purchased as indicated: Dulbecco modified Eagle’s medium(DMEM) supplemented with 4.5 g/liter glucose, L-glutamine, and sodium pyruvate (catalog number 10-013-CV; Corning, Inc.), minimum essential media (MEM) with L-glutamine (catalog number 10-010-CV; Corning, Inc.), Media 199 (catalog number 11150067, Thermo Scientific), OptiMEM media (catalog number 31985070, Life Technologies), FluoroBrite™ DMEM media (catalog number A18967-02, Life Technologies), trypsin/EDTA (catalog number 25-053-CI, Mediatech), fetal bovine serum (FBS) (catalog number S11150H; Atlanta Biologicals), a 100x solution of penicillin-streptomycin (catalog number 45000-652; VWR), HEPES (catalog number HOL06, Caisson Labs) camostat mesylate (catalog number SML0057; Sigma-Aldrich), E-64 (catalog number sc-201276; Santa Cruz Biotechnology), nafamostat mesylate (catalog number 14837; Cayman Chemical Company), apilimod (catalog number HY-14644;MedChem Express), dimethyl sulfoxide (DMSO) (catalog number 26855; Sigma-Aldrich), Toluene (catalog number 34866, Sigma-Aldrich), dichloromethane (catalog number 270997, Sigma-Aldrich), ethanol (catalog number TX89125172HU, VWR), round cover glass number 1.5 (25 mm, catalog number C8-1.5H-N; Cellvis), polydimethylsiloxane (PDMS) (Sylgard, catalog number DC4019862; Krayden), isopropyl alcohol (catalog number 9080-03/MK303108; VWR), potassium hydroxide (catalog number 484016; Sigma-Aldrich), TPCK treated trypsin (catalog number PI20233, Thermo Fischer), Aprotinin from bovine lung (catalog number A1153, Sigma-Aldrich), wheat germ agglutinin (WGA)-Alexa Fluor647 (catalog number W32466; Invitrogen), paraformaldehyde (catalog number P6148; Sigma-Aldrich), di-basic sodium phosphate (catalog number BP329-1; Thermo Fisher Scientific), monobasic potassium phosphate (catalog number P5379; Sigma-Aldrich), sodium chloride (catalog number SX0420-5; EMD Millipore), poly-D-lysine (catalog number P6403, Sigma-Aldrich), potassium chloride (catalog number P217-500; Thermo Fisher Scientific), Tris (catalog number T-400-500; Goldbio), EDTA (catalog number E5134; Sigma-Aldrich), sucrose (catalog number S0389;Sigma-Aldrich), fetal calf serum (FCS) (catalog number SH30073.03; HyClone), bovine serum albumin (GE Healthcare), Triton X-100 (catalog number 28314; Thermo Fisher), Hoechst DNA stain (catalog number62249; Thermo Fisher), L-glutamine (catalog number G7513; Sigma), Hoechst DNA dye (catalog numberH6021; Sigma-Aldrich), and Alexa Fluor 647fluorescently labeled goat anti-rabbit antibody (catalog number A32733; Thermo Fisher). IRDye 800CW Goat anti-rabbit IgG secondary antibody (catalog number 926-32211; Li-COR). Hydroxy Dynasore (Dynasore-OH) outsourced by the Kirchhausen laboratory, Hydroxy Dynasore (Dynasore-OH) (catalog number 5364, Tocris).

### Cells

Vero (ATCC CRL-1586), Caco-2 (ATCC HTB-37), Calu-3 (ATCC HTB-55) were purchased from ATCC. Vero TMPRSS2 cells ectopically expressing TMPRSS2 cells were a gift from Siyuan Ding (Washington University St. Louis) (*4*). Vero TMPRSS2* were as described in in (*5*) Vero, Vero TMPRSS2, Vero TMPRSS2* and Caco-2 cells, were grown in 10% CO_2_ and at 37°C in DMEM supplemented with 10% fetal bovine serum, 1% penicillin-streptomycin, and 25 mM HEPES, pH 7.4. Calu-3 cells were grown in 10% CO_2_ and at 37°C in DMEM supplemented with 10% (Kirchhausen laboratory) or 20% (Balistreri laboratory) fetal bovine serum, 1% penicillin-streptomycin, and 25 mM HEPES, pH 7.4. Vero, Vero TMPRSS2 and Caco-2 cells grown to ∼90% confluency were split at a ratio of 1:10 every 3 to 5 days. Vero TMPRSS2* cells grown to 70% confluency were split at a ratio of 1:10 every 3 days. Calu-3 cells at ∼90% confluency were split at a ratio of 1:3 every 5-8 days. MA104 cells (ATCC CRL-2378.1) were grown in 5% CO_2_ and at 37°C in medium 199 supplemented with 10% FBS and 1% penicillin-streptomycin and upon reaching ∼90% confluency were split at a ratio of 1:3 every 3 days. SVG-A cells were grown in 5% CO_2_ and at 37°C in MEM supplemented with 10% fetal bovine serum, 1% penicillin-streptomycin, and 25 mM HEPES, pH 7.4. HEK293T cells used for lentivirus production were grown in DMEM supplemented with 10% FBS and grown at 37°C and 5% CO_2_.

### Generation of VSV-SARS-CoV-2 chimeras

The generation of a replication competent recombinant VSV chimera expressing eGFP where the glycoprotein G was replaced with spike (S) protein Wuhan-Hu-1 strain (VSV-eGFP-SARS-CoV-2) has been described (*13*). Additional VSV recombinants expressing eGFP and SARS-CoV-2 spike variants were prepared as follows. Briefly, spike genes of Delta (B.1.617.2), Omicron (B.1.1.529) were truncated to remove the C-terminal 63 nucleotides and were then cloned into an infectious molecular clone of VSV-eGFP in place of the native VSV G gene. To generate VSV-eGFP-P-SARS-CoV-2 the open reading frame of the glycoprotein in plasmid pVSV1(+)-eGFP-P (PMID 16339901) was replaced by a codon-optimized version of the coding sequence of the Wuhan strain of the SARS-CoV-2 spike with the last 21 amino acid deleted.

To rescue the recombinant VSVs, BSRT7/5 cells were infected with vaccinia virus encoding the bacteriophage T7 RNA polymerase (vTF7-3) (PMID 3095828), as a source of transcriptase for expression of VSV N, P, L, and G, and the full-length antigenomic RNA of VSV from individual plasmids. Cell culture fluids were harvested ∼72 h post-infection, clarified by centrifugation (5 min and 1,000 × g), and the supernatant filtered through a 0.22-μm filter. Viruses were isolated by plaque assay on Vero TMPRSS2 cells. Individual plaque isolates of virus were amplified on MA104 cells at an MOI of 0.01 in Medium 199 containing 2% FBS and 20 mM HEPES pH 7.7 (Millipore Sigma) at 34°C. Viral supernatants were harvested upon extensive cytopathic effect and clarified of cell debris by centrifugation at 1,000 x g for 5 min. Viral stock RNA was extracted using RNeasy Mini kit (QIAGEN), and S was amplified using OneStep RT-PCR Kit (QIAGEN) and sequence verified by Sanger sequencing (GENEWIZ). Aliquots of sequence verified virus were maintained at −80°C.

### VSV-eGFP-SARS-CoV-2 infection assays

Infection assays for the Wuhan, Delta or Omicron VSV-eGFP-SARS-CoV-2 chimeras were done at a final ∼ 80% confluency of cells plated one day before the infection assay as previously described (*3*). Briefly, cells were incubated for one hour with the appropriate VSV-eGFP-SARS-CoV-2 chimera suspended in DMEM containing 25 mM HEPES at the indicated pH, at 37°C, and 10 %CO2. Cells were then rinsed three times and further incubated for 7 hours in the same medium, followed by chemical fixation and spinning disc confocal microscopy imaging (see below) to score extent of infection by visualization of eGFP expressing cells.

The effect of inhibitors on infection was determined in cells pre-incubated with medium containing the appropriate inhibitor for one hr, followed by its presence during both the inoculation period and during the 7-hr duration of the infection assay. A pH bypass experimental design was used to test the effect of pH on infection. Briefly, cells always kept at 37°C and 10% CO2 were incubated for 1 hour in DMEM containing 25 mM HEPES, pH 7.4 and 40 µM dynasore-OH), followed by incubation of the cells for another hour with the appropriate VSV-eGFP-SARS-CoV-2 variant using the same medium including dynasore-OH and the required acidic pH. Cells were then rinsed three times and incubated for another 7 hours in DMEM containing 25 mM HEPES, pH 7.4 and dynasore-OH.

At the end of this 7-hr incubation, medium was changed to PBS containing 5 µg/mL WGA-Alexa647 and cells incubated for 30 sec to highlight their surface, then chemically fixed with 4% PFA in PBS for 15 minutes at room temperature, rinsed 3x with PBS and imaged within 24 hours using a spinning-disk confocal microscope equipped with a Zeiss 40x oil NA=1.4 objective (1 pixel, 0.33 µm). Random fields, containing a Z-stack of 20 consecutive optical planes taken 1 µm apart (*6*) were obtained and successful infection scored if the cytosolic eGFP fluorescence signal from maximum Z-projections was 1.4 times higher than that of uninfected cells imaged using the same visualization conditions.

### SARS-CoV-2 infection assays

All experiments with SARS-CoV-2 were performed in biosafety level 3 (BSL3) facilities at the University of Helsinki with appropriate institutional permits. Virus samples were obtained under Helsinki University Hospital laboratory research permit 30 HUS/32/2018§16. Virus titers were determined by a plaque assay in Vero TMPRSS2* cells. Cells in DMEM, supplemented with 10% FBS, 2 mM l-glutamine, 1% penicillin-streptomycin, and 20 mM HEPES (pH 7.2), were seeded 48 h before treatment at a density of 15,000 cells per well in 96-well imaging plates (catalog number 6005182; PerkinElmer). Dynasore-OH (Tocris) at indicated concentration or DMSO control were added to cells 60 minutes before infection at an MOI of 0.5 PFU per cell with virus added at indicated pH. After 2 hours of infection, cells were washed removing unbound virus and dynasore-OH. Infections were carried for 16 hours at 37°C with 5% CO_2_. Cells were then fixed with 4% paraformaldehyde in PBS for 30 min at room temperature before being processed for immunodetection of viral N protein, automated fluorescence imaging, and image analysis. Briefly, viral NP was detected with an in-house-developed rabbit polyclonal antibody (Cantuti-Castelvetri), counterstained with Alexa Fluor 647-conjugated goat anti-rabbit secondary antibody, and nuclear staining was done using Hoechst DNA dye. Automated fluorescence imaging was done using a Molecular Devices Image-Xpress Nano high-content epifluorescence microscope equipped with a 20× objective and a 4.7-megapixel CMOS (complementary metal oxide semiconductor) camera (pixel size, 0.332 μm). Image analysis was performed with CellProfiler-3 software (www.cellprofiler.org). Automated detection of nuclei was performed using the Otsu algorithm inbuilt in the software. To automatically identify infected cells, an area surrounding each nucleus (5-pixel expansion of the nuclear area) was used to estimate the fluorescence intensity of the viral NP immunolabeled protein, using an intensity threshold such that <0.01% of positive cells were detected in noninfected wells.

### VSV-SARS-CoV-2 Atto 565 labeling

Stock solutions of VSV-SARS-CoV-2 and its variants at a concentration of ∼150 µg/ml viral RNA were conjugated with Atto 565-NHS ester (Sigma-Aldrich, cat. 72464) as previously described (*7*). Aliquots of Atto 565 NHS ester resuspended in anhydrous DMSO (Sigma) were dried down under house vacuum to a final yield of 0.25 µg/tube stocks and stored at -20°C. A sample of 100 µL of virus stock was mixed with 10 µL of 1 M NaHCO_3_ pH 8.3 and then added to one such tube containing dried Atto 565 NHS ester and incubated in the dark for 1 hour at room temperature; the reaction was ended by addition of 20 µL of 1 M Tris pH 8.0. Free Atto 565 dye was removed from labeled virus using a 0.5 mL Pierce desalting Zeba spin column of 40 kDa molecular weight cut off according to the manufacturer’s instructions, and labeled virus stored at 4°C in the dark was used within one week of labeling.

### VSV-SARS-CoV-2 Atto 565 labeling

Stock solutions of VSV-SARS-CoV-2 and its variants at a concentration of ∼150 µg/ml viral RNA were conjugated with Atto 565-NHS ester (Sigma-Aldrich, cat. 72464) as previously described (*7*). Aliquots of Atto 565 NHS ester resuspended in anhydrous DMSO (Sigma) were dried down under house vacuum to a final yield of 0.25 µg/tube stocks and stored at -20°C. A sample of 100 µL of virus stock was mixed with 10 µL of 1 M NaHCO_3_ pH 8.3 and then added to one such tube containing dried Atto 565 NHS ester and incubated in the dark for 1 hour at room temperature; the reaction was ended by addition of 20 µL of 1 M Tris pH 8.0. Free Atto 565 dye was removed from labeled virus using a 0.5 mL Pierce desalting Zeba spin column of 40 kDa molecular weight cut off according to the manufacturer’s instructions, and labeled virus stored at 4°C in the dark was used within one week of labeling.

### VSV-SARS-CoV-2 Atto 565 labeling

Stock solutions of VSV-SARS-CoV-2 and its variants at a concentration of ∼150 µg/ml viral RNA were conjugated with Atto 565-NHS ester (Sigma-Aldrich, cat. 72464) as previously described (*7*). Aliquots of Atto 565 NHS ester resuspended in anhydrous DMSO (Sigma) were dried down under house vacuum to a final yield of 0.25 µg/tube stocks and stored at -20°C. A sample of 100 µL of virus stock was mixed with 10 µL of 1 M NaHCO_3_ pH 8.3 and then added to one such tube containing dried Atto 565 NHS ester and incubated in the dark for 1 hour at room temperature; the reaction was ended by addition of 20 µL of 1 M Tris pH 8.0. Free Atto 565 dye was removed from labeled virus using a 0.5 mL Pierce desalting Zeba spin column of 40 kDa molecular weight cut off according to the manufacturer’s instructions, and labeled virus stored at 4°C in the dark was used within one week of labeling.

### Single molecule Atto 565 dye calibration

The number of Atto 565 molecules attached to a single virion was determined by comparing the total fluorescence intensities associated with a given virion and the fluorescence intensity associated with the last bleaching step of the same virion, as previously described (*1, 8*). This strategy corrects for local variations in the imaging conditions of the microscope and allows direct use of data from calibration results obtained i*n vitro* with the total fluorescence intensity acquired in cells REF. Briefly, fluorescently tagged VSV chimeras (0.1 µg/mL viral RNA) dissolved in PBS were adsorbed for 10 min to glass that had been extensively washed in 3x PBS after coated for 30 minutes and at room temperature with 0.1 mg/mL of poly-D-lysine. After removing unbound virus by 3 consecutive washes with PBS, the sample was imaged using spinning disc confocal microscopy using a 63x NA 1.4 oil objective or LLSM in scanning mode. Automated location of the fluorescent virions in the coverslip and corresponding fluorescence intensity determined by Gaussian fitting of images was done using the 3D CME package (*9*). To determine the number of Atto 565 dyes associated per each virus particle, the exposure was increased to one second and consecutive images acquired on the plane associated with the maximum original intensity. This continuous illumination protocol, conducent to extensive bleaching and ending with resolvable single bleaching steps, was used to determine the fluorescence of a single Atto 565 dye with the aid of a step detecting algorithm encoded in MATLAB (*10*).

### Western Blot analysis

Western blot analysis of VSV-SARS-CoV-2 samples were done as follows. 20 µL of virion sample at 10 µg/mL virus RNA was mixed with an equal volume of 2x concentrated Laemmli sample buffer, boiled for 10 minutes at 95°C and subjected to SDS-PAGE separation using Mini-PROTEAN any-kD 4-20% precast gels and electrically transferred to a nitrocellulose membrane. Transfer efficiency was verified using A ponceau staining. Nonspecific binding antibody to the membranes was prevented by incubation with 5% milk powder in TBS-T (50 mM TRIS, 150 mM NaCl, 0.1% Tween) for 1 hour a room temperature. The membranes were incubated overnight at 4°C with a solution containing 5% milk powder TBS-T and antibody specific for S used diluted 1 to 1000 from 1 µg/mL. Membranes were washed 3x with TBS-T, then incubated for one hour at room temperature with a 1 to 20,000 dilution of 1 mg/mL IRDye 800CW Goat anti-rabbit IgG secondary antibody in 5% milk powder TBS-T. Membranes were then washed 3x with what and scanned using a Licore Odyssey scanner.

### Preparation of glass coverslips

Infection and uptake assays of VSV-eGFP-SARS-CoV-2 and VSV-P-eGFP-SARS-CoV-2 done by spinning disc confocal microscopy visualization, were performed using 25 mm #1.5 coverslips bound with polydimethylsiloxane (PDMS) of about 1 mm in thickness and of 3 mm (infection) or 5 mm (uptake) in diameter wells punched as previously described (*9*). The punched PDMS sheets were attached onto the top surface of glass coverslips previously cleaned by sonication, first in isopropanol for 20 minutes and then in 0.5 M KOH for 20 minutes, followed by extensive washing in MilliQ water and ending with drying for 30 minutes at 90°C. The glass was then firmly pressed to the PDMS sheet, after exposure of their surfaces to air plasma in a PDC-001 plasma cleaner (Harrick Plasma) operating at 750 mtorr and 30W for 2 min. The bonded glass/PDMS was heated at 90°C for 20 min, followed by sterilization by incubation with 70% ethanol for 10 min immediately before plating of cells.

Single virus tracking experiments done using LLSM experiment were done using round #1.5 glass coverslips of 6 mm in diameter that had been cleaned by sequential 20-min steps of sonication, first in toluene, then in dichloromethane, followed by 100 % ethanol, ending in MilliQ water; the glass slides were stored in 70% ethanol until cells were plated. These glass coverslips were re-used at the culmination of the imaging experiments by following the same previous cleaning steps.

### Live cell spinning disc-confocal microscopy

Visualization experiments were done with an inverted spinning disc confocal microscope (*42*). Cells were plated one day prior to the experiment into 5 mm in diameter homemade PDMS wells to achieve 70% confluency during visualization. VSV-eGFP-P-SARS-CoV-2 was added to the well in DMEM containing 25 mM HEPES pH 7.4 for 1 hour. Cells where then labeled with 5 µg/mL of WGA-Alexa647 in FluoroBrite™ DMEM media containing 25 mM HEPES, pH 7.4 for 30 seconds followed by 3 rapid washes in the same media. Cells were then immediately imaged using a 63x objective on a spinning-disc confocal microscope taking the entire cell volume with 0.27 µm spacing between each optical plane. In parallel with the experiment in cells, VSV-eGFP-P-SARS-CoV-2 was bound to poly-D-lysine coated cover slips and imaged under the same

## REFERENCES

1. M. Hoffmann, H. Kleine-Weber, S. Schroeder, N. Krüger, T. Herrler, S. Erichsen, T. S. Schiergens, G. Herrler, N.-H. Wu, A. Nitsche, M. A. Müller, C. Drosten, S. Pöhlmann, SARS- CoV-2 Cell Entry Depends on ACE2 and TMPRSS2 and Is Blocked by a Clinically Proven Protease Inhibitor. Cell. 181, 271–280.e8 (2020).

2. D. Wrapp, N. Wang, K. S. Corbett, J. A. Goldsmith, C.-L. Hsieh, O. Abiona, B. S. Graham, J. S. McLellan, Cryo-EM structure of the 2019-nCoV spike in the prefusion conformation. Science. 367, 1260–1263 (2020).

3. J. Yang, S. J. L. Petitjean, M. Koehler, Q. Zhang, A. C. Dumitru, W. Chen, S. Derclaye, S. P. Vincent, P. Soumillion, D. Alsteens, Molecular interaction and inhibition of SARS-CoV-2 binding to the ACE2 receptor. Nat Commun. 11, 4541 (2020).

4. R. Yan, Y. Zhang, Y. Li, L. Xia, Y. Guo, Q. Zhou, Structural basis for the recognition of SARS-CoV-2 by full-length human ACE2. Sci New York N Y. 367, 1444–1448 (2020).

5. J. Shang, Y. Wan, C. Luo, G. Ye, Q. Geng, A. Auerbach, F. Li, Cell entry mechanisms of SARS-CoV-2. P Natl Acad Sci Usa. 117, 11727–11734 (2020).

6. C. B. Jackson, M. Farzan, B. Chen, H. Choe, Mechanisms of SARS-CoV-2 entry into cells. Nat Rev Mol Cell Biology. 23, 3–20 (2022).

7. B. A. Johnson, X. Xie, A. L. Bailey, B. Kalveram, K. G. Lokugamage, A. Muruato, J. Zou, X. Zhang, T. Juelich, J. K. Smith, L. Zhang, N. Bopp, C. Schindewolf, M. Vu, A. Vanderheiden, E. S. Winkler, D. Swetnam, J. A. Plante, P. Aguilar, K. S. Plante, V. Popov, B. Lee, S. C. Weaver, M. S. Suthar, A. L. Routh, P. Ren, Z. Ku, Z. An, K. Debbink, M. S. Diamond, P.-Y. Shi, A. N. Freiberg, V. D. Menachery, Loss of furin cleavage site attenuates SARS-CoV-2 pathogenesis. Nature. 591, 293–299 (2021).

8. R. Zang, M. F. G. Castro, B. T. McCune, Q. Zeng, P. W. Rothlauf, N. M. Sonnek, Z. Liu, K. F. Brulois, X. Wang, H. B. Greenberg, M. S. Diamond, M. A. Ciorba, S. P. J. Whelan, S. Ding, TMPRSS2 and TMPRSS4 promote SARS-CoV-2 infection of human small intestinal enterocytes. Sci Immunol. 5, eabc3582 (2020).

9. A. J. B. Kreutzberger, A. Sanyal, R. Ojha, J. D. Pyle, O. Vapalahti, G. Balistreri, T. Kirchhausen, Synergistic Block of SARS-CoV-2 Infection by Combined Drug Inhibition of the Host Entry Factors PIKfyve Kinase and TMPRSS2 Protease. J Virol. 95, e0097521 (2021).

10. X. Ou, Y. Liu, X. Lei, P. Li, D. Mi, L. Ren, L. Guo, R. Guo, T. Chen, J. Hu, Z. Xiang, Z. Mu, X. Chen, J. Chen, K. Hu, Q. Jin, J. Wang, Z. Qian, Characterization of spike glycoprotein of SARS-CoV-2 on virus entry and its immune cross-reactivity with SARS-CoV. Nat Commun. 11, 1620 (2020).

11. T. P. Peacock, D. H. Goldhill, J. Zhou, L. Baillon, R. Frise, O. C. Swann, R. Kugathasan, R. Penn, J. C. Brown, R. Y. Sanchez-David, L. Braga, M. K. Williamson, J. A. Hassard, E. Staller, B. Hanley, M. Osborn, M. Giacca, A. D. Davidson, D. A. Matthews, W. S. Barclay, The furin cleavage site in the SARS-CoV-2 spike protein is required for transmission in ferrets. Nat Microbiol. 6, 899–909 (2021).

12. D. Bestle, M. R. Heindl, H. Limburg, T. V. L. van, O. Pilgram, H. Moulton, D. A. Stein, K. Hardes, M. Eickmann, O. Dolnik, C. Rohde, H.-D. Klenk, W. Garten, T. Steinmetzer, E. Böttcher-Friebertshäuser, TMPRSS2 and furin are both essential for proteolytic activation of SARS-CoV-2 in human airway cells. Life Sci Alliance. 3, e202000786 (2020).

13. J. B. Case, P. W. Rothlauf, R. E. Chen, Z. Liu, H. Zhao, A. S. Kim, L.-M. Bloyet, Q. Zeng, S. Tahan, L. Droit, Ma. X. G. Ilagan, M. A. Tartell, G. Amarasinghe, J. P. Henderson, S. Miersch, M. Ustav, S. Sidhu, H. W. Virgin, D. Wang, S. Ding, D. Corti, E. S. Theel, D. H. Fremont, M. S. Diamond, S. P. J. Whelan, Neutralizing Antibody and Soluble ACE2 Inhibition of a Replication-Competent VSV-SARS-CoV-2 and a Clinical Isolate of SARS-CoV-2. Cell Host Microbe. 28, 475–485.e5 (2020).

14. B.-C. Chen, W. R. Legant, K. Wang, L. Shao, D. E. Milkie, M. W. Davidson, C. Janetopoulos, X. S. Wu, J. A. H. III, Z. Liu, B. P. English, Y. Mimori-Kiyosue, D. P. Romero, A. T. Ritter, J. Lippincott-Schwartz, L. Fritz-Laylin, R. D. Mullins, D. M. Mitchell, J. N. Bembenek, A.-C. Reymann, R. Böhme, S. W. Grill, J. T. Wang, G. Seydoux, U. S. Tulu, D. P. Kiehart, E. Betzig, Lattice light-sheet microscopy: Imaging molecules to embryos at high spatiotemporal resolution. Science. 346, 1257998 (2014).

15. C. J. Bradish, J. B. Kirkham, The Morphology of Vesicular Stomatitis Virus (Indiana C) Derived from Chick Embryos or Cultures of BHK21/13 Cells. Microbiology+. 44, 359–371 (1966).

16. D. K. Cureton, R. H. Massol, S. P. J. Whelan, T. Kirchhausen, The Length of Vesicular Stomatitis Virus Particles Dictates a Need for Actin Assembly during Clathrin-Dependent Endocytosis. Plos Pathog. 6, e1001127 (2010).

17. J. Klumperman, G. Raposo, The Complex Ultrastructure of the Endolysosomal System. Csh Perspect Biol. 6, a016857 (2014).

18. S. M. Ferguson, P. D. Camilli, Dynamin, a membrane-remodelling GTPase. Nat Rev Mol Cell Bio. 13, 75–88 (2012).

19. H. Damke, T. Baba, D. E. Warnock, S. L. Schmid, Induction of mutant dynamin specifically blocks endocytic coated vesicle formation. J Cell Biology. 127, 915–934 (1994).

20. E. Macia, M. Ehrlich, R. Massol, E. Boucrot, C. Brunner, T. Kirchhausen, Dynasore, a Cell-Permeable Inhibitor of Dynamin. Dev Cell. 10, 839–850 (2006).

21. Y.-L. Kang, Y. Chou, P. W. Rothlauf, Z. Liu, T. K. Soh, D. Cureton, J. B. Case, R. E. Chen, M. S. Diamond, S. P. J. Whelan, T. Kirchhausen, Inhibition of PIKfyve kinase prevents infection by Zaire ebolavirus and SARS-CoV-2. Proc National Acad Sci. 117, 20803–20813 (2020).

22. R. Zang, J. B. Case, E. Yutuc, X. Ma, S. Shen, M. F. G. Castro, Z. Liu, Q. Zeng, H. Zhao, J. Son, P. W. Rothlauf, A. J. B. Kreutzberger, G. Hou, H. Zhang, S. Bose, X. Wang, M. D. Vahey, K. Mani, W. J. Griffiths, T. Kirchhausen, D. H. Fremont, H. Guo, A. Diwan, Y. Wang, M. S. Diamond, S. P. J. Whelan, S. Ding, Cholesterol 25-hydroxylase suppresses SARS-CoV-2 replication by blocking membrane fusion. P Natl Acad Sci Usa. 117, 32105–32113 (2020).

23. S. Belouzard, V. C. Chu, G. R. Whittaker, Activation of the SARS coronavirus spike protein via sequential proteolytic cleavage at two distinct sites. Proc National Acad Sci. 106, 5871–5876 (2009).

24. Y. Liu, H.-Q. Qu, J. Qu, L. Tian, H. Hakonarson, Expression Pattern of the SARS-CoV-2 Entry Genes ACE2 and TMPRSS2 in the Respiratory Tract. Viruses. 12, 1174 (2020).

25. S. P. Sajuthi, P. DeFord, Y. Li, N. D. Jackson, M. T. Montgomery, J. L. Everman, C. L. Rios, E. Pruesse, J. D. Nolin, E. G. Plender, M. E. Wechsler, A. C. Y. Mak, C. Eng, S. Salazar, V. Medina, E. M. Wohlford, S. Huntsman, D. A. Nickerson, S. Germer, M. C. Zody, G. Abecasis, H. M. Kang, K. M. Rice, R. Kumar, S. Oh, J. Rodriguez-Santana, E. G. Burchard, M. A. Seibold, Type 2 and interferon inflammation regulate SARS-CoV-2 entry factor expression in the airway epithelium. Nat Commun. 11, 5139 (2020).

26. R. J. A. England, J. J. Homer, L. C. Knight, S. R. Ell, Nasal pH measurement: a reliable and repeatable parameter. Clin Otolaryngology Allied Sci. 24, 67–68 (1999).

27. B.-G. Kim, J.-H. Kim, S.-W. Kim, S.-W. Kim, K.-S. Jin, J.-H. Cho, J.-M. Kang, S.-Y. Park, Nasal pH in patients with chronic rhinosinusitis before and after endoscopic sinus surgery. Am J Otolaryng. 34, 505–507 (2013).

28. I. Glowacka, S. Bertram, M. A. Müller, P. Allen, E. Soilleux, S. Pfefferle, I. Steffen, T. S. Tsegaye, Y. He, K. Gnirss, D. Niemeyer, H. Schneider, C. Drosten, S. Pöhlmann, Evidence that TMPRSS2 Activates the Severe Acute Respiratory Syndrome Coronavirus Spike Protein for Membrane Fusion and Reduces Viral Control by the Humoral Immune Response. J Virol. 85, 4122–4134 (2011).

29. G. Simmons, S. Bertram, I. Glowacka, I. Steffen, C. Chaipan, J. Agudelo, K. Lu, A. J. Rennekamp, H. Hofmann, P. Bates, S. Pöhlmann, Different host cell proteases activate the SARS-coronavirus spike-protein for cell–cell and virus–cell fusion. Virology. 413, 265–274 (2011).

30. J. K. Millet, G. R. Whittaker, Host cell entry of Middle East respiratory syndrome coronavirus after two-step, furin-mediated activation of the spike protein. Proc National Acad Sci. 111, 15214–15219 (2014).

31. D. L. Floyd, J. R. Ragains, J. J. Skehel, S. C. Harrison, A. M. van Oijen, Single-particle kinetics of influenza virus membrane fusion. Proc National Acad Sci. 105, 15382–15387 (2008).

32. T. Ivanovic, J. L. Choi, S. P. Whelan, A. M. van Oijen, S. C. Harrison, Influenza-virus membrane fusion by cooperative fold-back of stochastically induced hemagglutinin intermediates. Elife. 2, e00333 (2013).

33. L. H. Chao, D. E. Klein, A. G. Schmidt, J. M. Peña, S. C. Harrison, Sequential conformational rearrangements in flavivirus membrane fusion. Elife. 3, e04389 (2014).

34. I. S. Kim, S. Jenni, M. L. Stanifer, E. Roth, S. P. J. Whelan, A. M. van Oijen, S. C. Harrison, Mechanism of membrane fusion induced by vesicular stomatitis virus G protein. Proc National Acad Sci. 114, E28–E36 (2017).

35. B. Meng, A. Abdullahi, I. A. T. M. Ferreira, N. Goonawardane, A. Saito, I. Kimura, D. Yamasoba, P. P. Gerber, S. Fatihi, S. Rathore, S. K. Zepeda, G. Papa, S. A. Kemp, T. Ikeda, M. Toyoda, T. S. Tan, J. Kuramochi, S. Mitsunaga, T. Ueno, K. Shirakawa, A. Takaori-Kondo, T. Brevini, D. L. Mallery, O. J. Charles, T. C.-N. B. C.-19 Collaboration, S. Baker, G. Dougan, C. Hess, N. Kingston, P. J. Lehner, P. A. Lyons, N. J. Matheson, W. H. Ouwehand, C. Saunders, C. Summers, J. E. D. Thaventhiran, M. Toshner, M. P. Weekes, P. Maxwell, A. Shaw, A. Bucke, J. Calder, L. Canna, J. Domingo, A. Elmer, S. Fuller, J. Harris, S. Hewitt, J. Kennet, S. Jose, J. Kourampa, A. Meadows, C. O’Brien, J. Price, C. Publico, R. Rastall, C. Ribeiro, J. Rowlands, V. Ruffolo, H. Tordesillas, B. Bullman, B. J. Dunmore, S. Gräf, J. Hodgson, C. Huang, K. Hunter, E. Jones, E. Legchenko, C. Matara, J. Martin, F. Mescia, C. O’Donnell, L. Pointon, J. Shih, R. Sutcliffe, T. Tilly, C. Treacy, Z. Tong, J. Wood, M. Wylot, A. Betancourt, G. Bower, C. Cossetti, A. D. Sa, M. Epping, S. Fawke, N. Gleadall, R. Grenfell, A. Hinch, S. Jackson, I. Jarvis, B. Krishna, F. Nice, O. Omarjee, M. Perera, M. Potts, N. Richoz, V. Romashova, L. Stefanucci, M. Strezlecki, L. Turner, E. M. D. D. D. Bie, K. Bunclark, M. Josipovic, M. Mackay, H. Butcher, D. Caputo, M. Chandler, P. Chinnery, D. Clapham-Riley, E. Dewhurst, C. Fernandez, A. Furlong, Graves, J. Gray, S. Hein, T. Ivers, E. L. Gresley, R. Linger, M. Kasanicki, R. King, S. Meloy, A. Moulton, F. Muldoon, N. Ovington, S. Papadia, C. J. Penkett, I. Phelan, V. Ranganath, R. Paraschiv, A. Sage, J. Sambrook, I. Scholtes, K. Schon, H. Stark, K. E. Stirrups, P. Townsend, N. Walker, J. Webster, T. G. to P. J. (G2P-J. Consortium, E. P. Butlertanaka, Y. L. Tanaka, J. Ito, K. Uriu, Y. Kosugi, M. Suganami, A. Oide, M. Yokoyama, M. Chiba, C. Motozono, H. Nasser, R. Shimizu, K. Kitazato, H. Hasebe, T. Irie, S. Nakagawa, J. Wu, M. Takahashi, T. Fukuhara, K. Shimizu, K. Tsushima, H. Kubo, Y. Kazuma, R. Nomura, Y. Horisawa, K. Nagata, Y. Kawai, Y. Yanagida, Y. Tashiro, K. Tokunaga, S. Ozono, R. Kawabata, N. Morizako, K. Sadamasu, H. Asakura, M. Nagashima, K. Yoshimura, E.-C. Consortium, P. Cárdenas, E. Muñoz, V. Barragan, S. Márquez, B. Prado-Vivar, M. Becerra-Wong, M. Caravajal, G. Trueba, P. Rojas-Silva, M. Grunauer, B. Gutierrez, J. J. Guadalupe, J. C. Fernández-Cadena, D. Andrade-Molina, M. Baldeon, A. Pinos, J. E. Bowen, A. Joshi, A. C. Walls, L. Jackson, D. Martin, K. G. C. Smith, J. Bradley, J. A. G. Briggs, J. Choi, E. Madissoon, K. B. Meyer, P. Mlcochova, L. Ceron-Gutierrez, R. Doffinger, S. A. Teichmann, A. J. Fisher, M. S. Pizzuto, A. de Marco, D. Corti, M. Hosmillo, J. H. Lee, L. C. James, L. Thukral, D. Veesler, A. Sigal, F. Sampaziotis, I. G. Goodfellow, K. Sato, R. K. Gupta, Altered TMPRSS2 usage by SARS-CoV-2 Omicron impacts infectivity and fusogenicity. Nature. 603, 706–714 (2022).

36. H. Zhao, L. Lu, Z. Peng, L.-L. Chen, X. Meng, C. Zhang, J. D. Ip, W.-M. Chan, A. W.-H. Chu, K.-H. Chan, D.-Y. Jin, H. Chen, K.-Y. Yuen, K. K.-W. To, SARS-CoV-2 Omicron variant shows less efficient replication and fusion activity when compared with Delta variant in TMPRSS2-expressed cells. Emerg Microbes Infec. 11, 277–283 (2022).

37. D. Bojkova, M. Widera, S. Ciesek, M. N. Wass, M. Michaelis, J. Cinatl, Reduced interferon antagonism but similar drug sensitivity in Omicron variant compared to Delta variant of SARS-CoV-2 isolates. Cell Res. 32, 319–321 (2022).

38. R. M. Effros, F. P. Chinard, The in vivo pH of the extravascular space of the lung. J Clin Invest. 48, 1983–1996 (1969).

39. R. D. de Vries, K. S. Schmitz, F. T. Bovier, C. Predella, J. Khao, D. Noack, B. L. Haagmans, S. Herfst, K. N. Stearns, J. Drew-Bear, S. Biswas, B. Rockx, G. McGill, N. V. Dorrello, S. H. Gellman, C. A. Alabi, R. L. de Swart, A. Moscona, M. Porotto, Intranasal fusion inhibitory lipopeptide prevents direct-contact SARS-CoV-2 transmission in ferrets. Sci New York N Y. 371, 1379–1382 (2021).

40. F. Aguet, S. Upadhyayula, R. Gaudin, Y. Chou, E. Cocucci, K. He, B.-C. Chen, K. Mosaliganti, M. Pasham, W. Skillern, W. R. Legant, T.-L. Liu, G. Findlay, E. Marino, G. Danuser, S. Megason, E. Betzig, T. Kirchhausen, Membrane dynamics of dividing cells imaged by lattice light-sheet microscopy. Mol Biol Cell. 27, 3418–3435 (2016).

41. A. H. Abdelhakim, E. N. Salgado, X. Fu, M. Pasham, D. Nicastro, T. Kirchhausen, S. C. Harrison, Structural Correlates of Rotavirus Cell Entry. Plos Pathog. 10, e1004355 (2014).

42. E. Cocucci, F. Aguet, S. Boulant, T. Kirchhausen, The First Five Seconds in the Life of a Clathrin-Coated Pit. Cell. 150, 495–507 (2012).

43. Y.-Y. Chou, S. Upadhyayula, J. Houser, K. He, W. Skillern, G. Scanavachi, S. Dang, A. Sanyal, K. G. Ohashi, G. D. Caprio, A. J. B. Kreutzberger, T. J. Vadakkan, T. Kirchhausen, Inherited nuclear pore substructures template post-mitotic pore assembly. Dev Cell. 56, 1786–1803.e9 (2021).

44. Y. Chen, N. C. Deffenbaugh, C. T. Anderson, W. O. Hancock, Molecular counting by photobleaching in protein complexes with many subunits: best practices and application to the cellulose synthesis complex. Mol Biol Cell. 25, 3630–3642 (2014).

## REFERENCES

1. Y.-Y. Chou, S. Upadhyayula, J. Houser, K. He, W. Skillern, G. Scanavachi, S. Dang, A. Sanyal, K. G. Ohashi, G. D. Caprio, A. J. B. Kreutzberger, T. J. Vadakkan, T. Kirchhausen, Inherited nuclear pore substructures template post-mitotic pore assembly. Dev Cell. 56, 1786–1803.e9 (2021).

2. Y.-L. Kang, Y. Chou, P. W. Rothlauf, Z. Liu, T. K. Soh, D. Cureton, J. B. Case, R. E. Chen, M. S. Diamond, S. P. J. Whelan, T. Kirchhausen, Inhibition of PIKfyve kinase prevents infection by Zaire ebolavirus and SARS-CoV-2. Proc National Acad Sci. 117, 20803–20813 (2020).

3. A. J. B. Kreutzberger, A. Sanyal, R. Ojha, J. D. Pyle, O. Vapalahti, G. Balistreri, T. Kirchhausen, Synergistic Block of SARS-CoV-2 Infection by Combined Drug Inhibition of the Host Entry Factors PIKfyve Kinase and TMPRSS2 Protease. J Virol. 95, e0097521 (2021).

4. R. Zang, M. F. G. Castro, B. T. McCune, Q. Zeng, P. W. Rothlauf, N. M. Sonnek, Z. Liu, K. F. Brulois, X. Wang, H. B. Greenberg, M. S. Diamond, M. A. Ciorba, S. P. J. Whelan, S. Ding, TMPRSS2 and TMPRSS4 promote SARS-CoV-2 infection of human small intestinal enterocytes. Sci Immunol. 5, eabc3582 (2020).

5. L. Cantuti-Castelvetri, R. Ojha, L. D. Pedro, M. Djannatian, J. Franz, S. Kuivanen, F. van der Meer, K. Kallio, T. Kaya, M. Anastasina, T. Smura, L. Levanov, L. Szirovicza, A. Tobi, H. Kallio-Kokko, P. Österlund, M. Joensuu, F. A. Meunier, S. J. Butcher, M. S. Winkler, B. Mollenhauer, A. Helenius, O. Gokce, T. Teesalu, J. Hepojoki, O. Vapalahti, C. Stadelmann, G. Balistreri, M. Simons, Neuropilin-1 facilitates SARS-CoV-2 cell entry and infectivity. Sci New York N Y. 370, 856–860 (2020).

6. E. Cocucci, F. Aguet, S. Boulant, T. Kirchhausen, The First Five Seconds in the Life of a Clathrin-Coated Pit. Cell. 150, 495–507 (2012).

7. A. H. Abdelhakim, E. N. Salgado, X. Fu, M. Pasham, D. Nicastro, T. Kirchhausen, S. C. Harrison, Structural Correlates of Rotavirus Cell Entry. Plos Pathog. 10, e1004355 (2014).

8. E. Cocucci, F. Aguet, S. Boulant, T. Kirchhausen, The First Five Seconds in the Life of a Clathrin-Coated Pit. Cell. 150, 495–507 (2012).

9. F. Aguet, S. Upadhyayula, R. Gaudin, Y. Chou, E. Cocucci, K. He, B.-C. Chen, K. Mosaliganti, M. Pasham, W. Skillern, W. R. Legant, T.-L. Liu, G. Findlay, E. Marino, G. Danuser, S. Megason, E. Betzig, T. Kirchhausen, Membrane dynamics of dividing cells imaged by lattice light-sheet microscopy. Mol Biol Cell. 27, 3418–3435 (2016).

10. Y. Chen, N. C. Deffenbaugh, C. T. Anderson, W. O. Hancock, Molecular counting by photobleaching in protein complexes with many subunits: best practices and application to the cellulose synthesis complex. Mol Biol Cell. 25, 3630–3642 (2014).

